# Fast photoswitchable molecular prosthetics control neuronal activity in the cochlea

**DOI:** 10.1101/2021.05.25.445123

**Authors:** Aida Garrido-Charles, Antoine Huet, Carlo Matera, Anupriya Thirumalai, Amadeu Llebaria, Tobias Moser, Pau Gorostiza

## Abstract

Artificial control of neuronal activity enables studies of neural circuits and restoration of neural function. Direct, rapid, and sustained photocontrol of intact neurons could overcome shortcomings of established electrical stimulation such as poor selectivity. We have developed fast photoswitchable ligands of glutamate receptors to establish such control in the auditory system. The new photoswitchable ligands produced photocurrents in untransfected neurons upon covalently tethering to endogenous glutamate receptors and activating them reversibly with visible light pulses of few milliseconds. As a proof of concept of these molecular prostheses, we apply them to the ultrafast synapses of auditory neurons of the cochlea that encode sound and provide auditory input to the brain. This drug-based method affords kilohertz rate stimulation of auditory neurons of adult gerbils without genetic manipulation that would be required for their optogenetic control. The new photoswitchable ligands are also broadly applicable to spatiotemporally control fast spiking interneurons in the brain.

## INTRODUCTION

Ultrafast signaling is a feature of several important neural circuits such as in inner ear, the brainstem, the cerebellum and the cerebrum (Kaczmarek & Zhang, 2017; Hu *et al*, 2014; Grothe *et al*, 2010). Such signaling builds on specialized synapses for synchronous neurotransmission as well as on suitable neural membrane properties for action potential generation and propagation. The resulting neural firing features high rates and submillisecond precision. For an example, synaptic sound encoding builds on ultrafast glutamatergic transmission at specialized ribbon synapses achieving firing at rates of several hundreds of Hertz with submillisecond precision (Huet *et al*, 2016; Moser *et al*, 2019). Utmost precision of the neural time code of the incoming sound forms the basis of sound localization in dedicated neural circuits of the brainstem that feature powerful calyceal synapses and neurons with extremely short membrane time constants owing to their specialized set of ion channels (Kaczmarek & Zhang, 2017; Grothe *et al*, 2010).

Dissecting the function of such time-critical neural circuitries requires ultrafast control of neuronal activity. Likewise, functional restoration, for example, following degeneration of the sensory receptor cells, needs approaches that re-instate the physiological behavior as closely as possible. Restoration of hearing to the deaf currently employs electrical stimulation of spiral ganglion neurons (SGNs) by cochlear implants (CI, (Kleinlogel *et al*, 2020; Zeng, 2017; Lenarz, 2017). Due to wide spread of current from each electrode contact encoding of sound frequency information is heavily limited in CIs. Moreover, electrical SGN stimulation results in supernatural temporal precision of spiking and hence CIs employ high rates to generate pseudostochasticity of SGN firing. Recently, optogenetics has been proposed for improved bionic SGN stimulation, as light can be better confined in space and optogenetically-evoked firing shows near-physiological temporal fidelity. Yet, it requires genetic manipulation for expression of channelrhodopsins. Avoiding the need of gene therapy, photopharmacology could help reducing the complexity of optical SGN stimulation. Reversibility of chemical photoswitches (Ankenbruck *et al*, 2018) make them interesting candidates for controlling neural excitation via endogenous neuronal receptors, such as ionotropic glutamate receptors of the postsynaptic SGN boutons in the cochlea.

Photoswitchable tethered ligands (PTLs) which are covalently attached to their receptor seem particularly attractive for this purpose as they provide more precise photocontrol than freely diffusible photochromic ligands (PCLs). PTLs produce higher local concentrations (Gorostiza *et al*, 2007) and cannot diffuse away, which yields a sharper separation in biological activity between the two isomeric states (Hüll *et al*, 2018; Reiner & Isacoff, 2014). Genetic manipulation can be avoided with affinity labeling conjugation of PTLs to target native nucleophilic residues in the protein (e.g. lysine (Izquierdo-Serra *et al*, 2016), histidine (Harvey & Trauner, 2008)). These targeted covalent photoswitches (TCPs) can be applied to intact neurons and readily provide photocontrol for hours (Izquierdo-Serra *et al*, 2016). However, their bistable i.e. dual-color, slow-relaxing photochromism (Volgraf *et al*, 2006; Gorostiza & Isacoff, 2007, 2008; Berlin *et al*, 2016; Kienzler & Isacoff, 2017) has so far hindered their application to ultrafast synapses. Thus, methods for direct, rapid and sustained photocontrol of activity in intact neurons constitute an unmet need, both to study neuronal circuits for basic research purposes and to explore new phototherapies. Interestingly, when sensory neurons are damaged or absent, fast and sustained neurotransmitter release is impaired, but the postsynaptic neurons and receptors retain their full capacities (activation kinetics, localization, complex formation with regulatory proteins) for extended periods of time, offering untapped potential for functional restoration (Kleinlogel *et al*, 2020).

Here, we developed a fast-switching glutamate TCP (TCP_fast_, Figure 1.A) that fulfills the above-mentioned requirements. We show the ability of TCP_fast_ to produce photocurrents in naive hippocampal neurons likely via tethering to endogenous glutamate receptors and reversibly modulating their activity with visible light pulses as short as a few milliseconds. As an original proof of concept, we demonstrate the application of this molecular tool to the ultrafast first synapses of the auditory system formed by inner hair cells (IHCs) and spiral ganglion neurons (SGNs), which mediate cochlear sound encoding. This drug-based method affords kilohertz rate stimulation in cochlear SGNs of wildtype adult gerbils, matching the performance of optogenetic photostimulation that requires gene therapy. In general, these photoswitches provide a fundamental resource of broad interest to spatiotemporally control endogenous receptors in intact neuronal circuits with ultrafast signaling.

**Figure 1.**
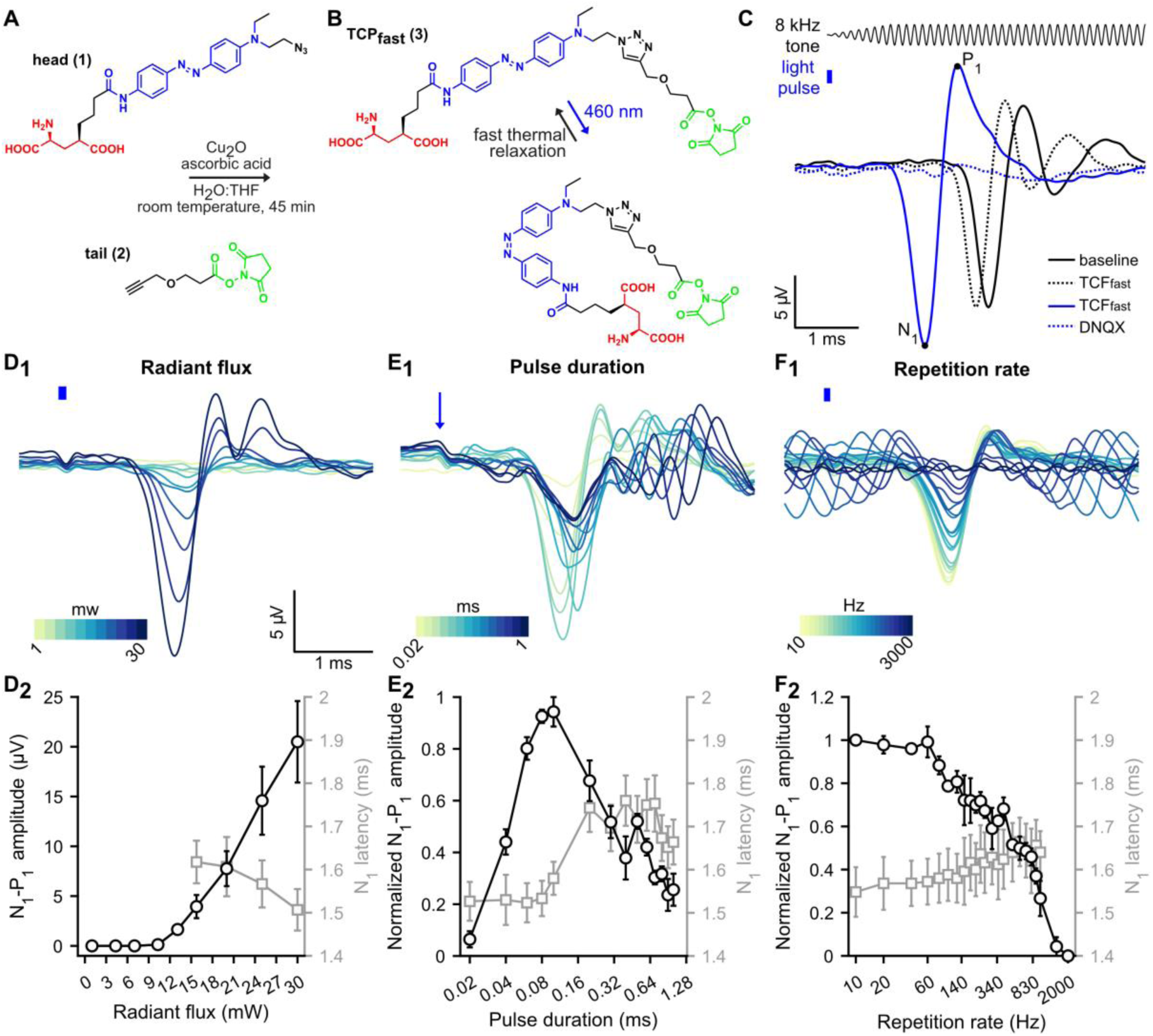
Fast photoswitchable control of SGN activity mediated by TCP_fast_. **A.** Molecular design showing ’head’ and ’tail’ precursors of TCP_fast_ that are freshly coupled prior to incubation in neuronal tissue. **B.** Photoisomerization between the *cis* (blue light, λ = 460 nm) and *trans* (dark, fast relaxation) conformations. **C.** Representative acoustically- (black, aCAP) and optically-evoked CAP (blue, oCAP) following 2.5 µM application of TCP_fast_. aCAP (black, 200 averages, 8 kHz toneburst, 30 dB SPL, repetition rate = 20 Hz) are similar before and after TCP_fast_ application. oCAP (blue, 80 µs, 30 mW, repetition rate = 10 Hz) is abolished following application of competitive antagonist DNQX (1 mM). Stimuli are represented on the top. **D_1_-F_1_**. Representative oCAP in response to various radiant fluxes (80 μs at 10 Hz, D_1_), pulse durations (27 mW at 10 Hz, E_1_) and repetition rates (80 μs at 27 mW, F_1_). In D_1_ and F_1_, the blue line indicates the light stimuli and in E_1_ the blue arrow indicates the beginning of the light pulse. A color scale is used to represent the variable. **D_2_-F_2_**. Quantification (*n* = 6 cochleae) of the oCAP amplitude (N_1_-P_1_, black axis) and oCAP latency (N_1_, gray axis) as a function of the radiant flux (D_2_), the pulse duration (E_2_) and the repetition rate (F_2_). In E_2_ and F_2_, oCAP amplitudes were expressed as relative to the highest amplitude recorded for the given measure.

## RESULTS

### Design and synthesis of a fast-switching TCP of ionotropic glutamate receptors

The molecular design of TCP_fast_ (**3**) was based on the recently reported TCPs (Izquierdo-Serra *et al*, 2016) and is shown in Figure 1.A. TCPs have a modular structure obtained by combining a ‘head’, which bears both the bioactive ligand (glutamate moiety) and the photoisomerizable unit (azobenzene) (Supplementary Scheme S1), and a ‘tail’ bearing the anchoring group (NHS ester). NHS ester-activated linkers are short-lived groups that promptly react with primary amines (e.g., lysine residues) in neutral or slightly alkaline conditions (pH 7.2−9). To avert self-reactivity, TCPs are readily generated prior to attachment to the target protein via a copper(I)-catalyzed azide-alkyne cycloaddition reaction (CuAAC, also known as “click chemistry”) (Supplementary Scheme S2).

Previous TCPs featured azobenzene moieties that were photoisomerized using two different illumination wavelengths (380 and 500 nm) and characterized by slow thermal *cis*-to-*trans* relaxation (Izquierdo-Serra *et al*, 2016). Azobenzenes displaying faster relaxation kinetics and single, longer wavelength switching can be obtained with minimal variation of their chemical structure by generating a “push–pull” system. It consists of including electron donating groups on one side of the azo unit and electron withdrawing groups on the other to lower the energy barrier of the *cis*-to-*trans* isomerization (Chi *et al*, 2006; Bandara & Burdette, 2012). This also results in a red-shifting of the azobenzene absorption spectrum, which is useful to reduce light scattering and the potential phototoxicity, which is greater for violet light, for *in vivo* application. Thus, we designed a TCP_fast_ head in which one of the two amide groups at the para positions of the azobenzene was replaced by a tertiary amine as electron-donating group (Kienzler *et al*, 2013). To avoid perturbing the ligand region, we chose to introduce this modification on the opposite side of the azobenzene core (head compound **1**). Compound **1** was prepared via a 5-step synthesis starting from commercially available materials (Supplementary Scheme S1 and SI for details). This photoswitch showed an absorption maximum at about 460 nm (blue light) in aqueous solution at neutral pH, as previously reported (see SI). Moreover, no variation of the absorption spectrum could be detected by steady-state UV-Vis spectroscopy, suggesting that it rapidly (<1 s) relaxes back to *trans* when the light is turned off. The head (**1**) was coupled via CuAAC with a commercial tail (compound **2**) providing a fast-relaxing and red-shifted ligand (**3**) with similar length to TCP9 (Izquierdo-Serra *et al*, 2016) (Figure 1.B and Supplementary Scheme S2). In order to reach satisfactory head-tail coupling rate in minutes at room temperature, we replaced the conventional sodium ascorbate by ascorbic acid (see SI for details). We hypothesized that the buffering effect of the ascorbic acid (to the tertiary amine in compound **1**) could create more proper conditions to promote the catalytic cycle of the reaction (Shao *et al*, 2010) as well as favor the formation of active copper(I) species from the copper(I) oxide catalyst (Shao *et al*, 2011). Since NHS-based ligands are constitutively short-lived, we confirmed the formation of the desired TCP_fast_ by liquid chromatography–mass spectrometry (LC-MS) of the click reaction crudes and verified their ability to conjugate primary amine-containing biomolecules by reacting them with pure lysine as a mock protein residue (see SI for details).

### Characterization of TCP_fast_ in cultured neurons

We made use of the several glutamate receptors (GluR) subunits expressed by hippocampal neurons (Janssens & Lesage, 2001) to evaluate, on dissociated neuron cultures, the ability of TCP_fast_ to photocontrol glutamate receptor activity. TCP_fast_ was conjugated to GluRs by using the same incubation conditions (i.e. 2 min at 25-100 µM, pH 9 to favor deprotonation and reactivity of nucleophilic residues in the receptors followed by wash-out of physiological solution, pH 7.4) previously shown to be favorable for TCP conjugation (Izquierdo-Serra *et al*, 2016). Using whole cell patch clamp recordings, 473 nm illumination did not elicit photocurrent directly after incubation with TCP_fast_. However, in the additional presence of glutamate (300 µM), 473 nm illumination elicited 2.5-220 pA photocurrents (Supplementary Figure S13). As intended by the chemical design, the *cis* isomer of TCP_fast_, induced by blue light illumination, evoked inward photocurrents thus supporting the fact that TCP_fast_ enabled photomodulation of glutamate receptors. Next, we tested the effect of light in absence of TCP_fast_ incubation (Supplementary Figure S14). The current measured at the onset of 300 µM glutamate perfusion were of similar amplitude in both cases (793±131 pA with TCP_fast_ and 1161±445 pA without TCP_fast_), discarding any non-specific effect of light stimulation.

Next, we showed that the number of photosensitized hippocampal neurons and the amplitude of the photocurrent evoked by blue light (Supplementary Figure S15) increased with TCP_fast_ concentration during the incubation (25, 50, 75 and 100 µM). At 25 µM of TCP_fast_, 47.62% of neurons (n = 21) had a measurable photocurrent (on average: 25 ± 12 pA) while 100 µM raised this to 90% of the neurons (n = 11) with a photocurrent of 93 ± 25 pA. In contrast, the TCP_fast_ relaxation lifetime in the dark seemed to be independent of TCP_fast_ concentration, amounted to 220 ± 48 ms (Supplementary Figure S16) and was, in accordance with our chemical design, faster than shown for TCPs (∼80 min) (Izquierdo-Serra *et al*, 2016). For a subset of neurons, we also found that the photocurrent increased in amplitude with the radiant flux (Supplementary Figure S17) and could be evoked by light pulse as short as 3 ms (Supplementary Figure S18).

Finally, we showed that the photoresponses can be reversibly blocked by the AMPA receptor antagonist DNQX (100 µM) but not by the NMDA receptor antagonist AP5 (100 µM) (Supplementary Figure S19). This demonstrates that TCP_fast_ photocurrents are receptor-specific and that the photoswitchable ligands are indeed covalently tethered to the GluRs (as they are not washed away by the competitive antagonists).

### *In vivo* photocontrol of neural activity in gerbil’s cochlea

The potential of TCP_fast_ to *in vivo* photosensitize SGNs was tested in the Mongolian gerbil cochlea using the same batch in the same click reaction condition than for *in vitro* characterization above. To do so, we applied TCP_fast_ to the niche of the round window (RW, i.e. one of the two openings into the cochlea) for passive diffusion into the cochlea and recorded acoustically and optically evoked cochlear mass potentials (Supplementary Figure S20.A_1-4_). Following a 10 min application of 12.5 µM TCP_fast_ (0.25% of organic solvent) and wash-out, the acoustically evoked potentials reflecting the outer hair cells (i.e. the cochlear microphonic, Supplementary Figure S20.A_3_ and S21.B), inner hair cells (i.e. the summating potential, Supplementary Figure S20.A_4_ and S20.C), and the SGNs (i.e. the compound action potential, CAP, Supplementary Figure S20.A_4_ and S20.D) remained largely unchanged (*n* = 4, Friedman’s test), indicating that TCP_fast_ incubation does not interfere with cochlear physiology. Upon subsequent optical fiber-based stimulation (blue laser, λ = 473 nm), we recorded transient optically evoked CAPs (oCAPs) in 12 out of 12 cochleae treated with TCP_fast_. oCAPs were similar in shape to acoustically evoked CAP and had an amplitude of 40.88 ± 12.58 µV (equivalent to the amplitude of a 50 dB SPL toneburst, Supplementary Figure S20.E). oCAPs vanished within minutes of optical stimulation, which was accompanied of a disappearance of the acoustic potentials reflecting the IHC and SGNs activation. The number of synapses per IHC (i.e. the juxtaposition of pre- [CtBP2] and post-synaptic [Homer 1] markers) was similar between treated and non-treated cochleae, at all frequencies, arguing against glutamate excitotoxicity induced by TCP_fast_–mediated optical stimulation (Supplementary Figure S20.F-G).

Next, we employed a lower TCP_fast_ dose (2.5 µM, <0.05 % of organic solvent) which enabled stable acoustically and optically evoked mass potentials in 60% of the incubated cochleae (Figure 1.C-F, Supplementary Figure S21.A-C. In 30% of the incubated cochleae, oCAPs could be recorded only transiently and loss of oCAP was accompanied by the light-induced loss of the acoustic potentials that reflect the activation of IHC and SGNs, Supplementary Figure S21.D-F). oCAPs were similar in shape and amplitude to aCAP evoked by a 40 dB SPL toneburst but consistent with direct neural excitation showed a shorter latency (∼1.5 ms, Figure 1.C). oCAPs could be abolished by DNQX (1 mM) application in the RW, confirming a response mediated by AMPA receptors (Figure 1.C, dashed blue trace).

We further characterized the TCP_fast_-mediated optical response by measuring oCAP amplitude and latency as a function of radiant flux (*n* = 6) using 80 µs light pulses at a repetition rate of 10 Hz (Figure 1.D_1-2_). oCAP threshold amounted to 12.5 ± 0.62 mW (1 ± 0.05 µJ). From there, oCAP amplitude increased linearly with radiant flux up to 20.5 ± 4.09 µV at 30 mW and latency decreased from 1.62 ± 0.05 ms to 1.51 ± 0.05 ms. Next, we measured the effect of the pulse duration on the oCAP (Figure 1.E_1-2_, 30 mW, repetition rate = 10 Hz): the biggest oCAPs were recorded in response to 80 and 100 µs light pulse. From there, oCAPs decreased in amplitude for shorter and longer durations. In 50% of the cases, sizable oCAP (i.e. > 1µV) were measured in light pulses as short as 20 µs, corresponding to 0.6 µJ. In response to light pulses shorter than 100 µs, oCAP were characterized by a single negative wave occurring at ∼1.54 ms. In contrast, oCAPs evoked by longer pulse had multiple negative peaks which might reflect the firing of multiple synchronous action potential across SGNs in response to the longer stimuli. Finally, we measured the effect of the repetition rate on oCAPs using 80 µs light pulses (Figure 1.F_1-2_, 30 mW). The oCAPs were stable in amplitude up to 60 Hz and decreased exponentially at higher repetition rate until not being observable above 1.5 kHz. For one cochlea for which DNQX (1 mM) application at the RW (Supplementary Figure S22) could be performed, oCAPs were initially observable up to 2 kHz and abolished by the DNQX, supporting the concept of fast control of AMPA receptor with TCP_fast_ in the SGNs.

## DISCUSSION

In this work, we developed a chemical-biological method, called TCP_fast_, which allows fast photoswitching of neuronal activity in native neurons. TCP_fast_ is a photoswitchable ligand that is chemically attached to endogenous receptors at the postsynaptic side (Izquierdo-Serra *et al*, 2016). The molecular design of TCP_fast_ optimized the properties of the bistable photoisomerizable group of former TCPs (Izquierdo-Serra *et al*, 2016), i.e. thermal relaxation lifetime and absorption wavelength, without altering the ligand and the reactive group. TCP_fast_ functions as a molecular prosthesis that bypasses the neurotransmitter-encoded signal by a photonic signal. Photosensitization of cochlear SGNs by locally administered TCP_fast_ enabled temporally precise light-evoked SGN firing up to a rate of 1.5 kHz, hence beyond the limits of the fastest optogenetic SGN stimulation (≤ 1kHz (Keppeler *et al*, 2018)). Hence, TCP_fast_-mediated photopharmacology might serve as an interesting alternative to the optogenetic approach (Dieter *et al*, 2020; Kleinlogel *et al*, 2020) for the development of an optical cochlear implant.

TCP_fast_ elicited depolarizing photocurrents that are activated by the *cis* isomer (‘*cis*-on’, which is generally preferred to avoid activation in the dark) and that are reversed in the absence of illumination with a thermal relaxation lifetime of 220 ms (Supplementary Figure S16). We can currently only speculate on how such thermal relaxation rate yields the ultrafast photostimulation of SGNs eliciting SGN CAPs with interstimulus intervals ranging from few milliseconds to sub-millisecond. Of note, postsynaptic boutons of SGNs likely contain hundreds of AMPAR such that each light stimulus might hit a sufficient number of AMPARs with the TCP_fast_ being in the *trans* state. Moreover, CAPs represent population responses and so light pulses might be recruiting variable subsets of SGNs. Future work could consider to further speed-up thermal relaxation potentially by replacing the current azobenzene unit in TCP_fast_ with faster photoswitches such as (hydroxy-substituted) phenylazopyrimidine (milliseconds to nanoseconds range (Camarero *et al*, 2020; Čechová *et al*, 2020; Garcia-Amorós *et al*, 2012)). However, such accelerated back-switching might in turn result in lower %*cis* conversion in the photostationary state and, therefore, require higher light intensity. In addition, further increasing the wavelength for photoswitching (Rullo *et al*, 2014) might improve the overall utility of the method due to better tissue penetration and lower phototoxicity risk.

We demonstrated, in dissociated hippocampal neurons, specific photomodulation of AMPA/kainate receptors by TCP_fast_. In this experimental setting, TCP_fast_ photomodulation required presence of glutamate in the bath solution, which we could not avoid at any of the concentration tested (Supplementary Figure S15). One may speculate that the requirement of glutamate could be caused by incomplete conjugation of TCP_fast_ to a heterogeneous population of endogenous GluRs. This concept may be translated into a distinctive subunit occupancy resulting in differential agonist affinity and efficacy and so reducing the channel open probability (gating) (Prieto & Wollmuth, 2010; Reiner & Isacoff, 2014; Jin *et al*, 2003). Another possible mechanism would be the presence of native GluR heterotetramers (Greger *et al*, 2017) with subunits displaying different TCP conjugation and photoswitching properties (Reiner et al, 2014).

Considering the unique properties of TCP_fast_ to modulate native GluR activity, TCP_fast_ was tested *in vivo* at the synapse between IHC and SGNs of the cochlea. TCP_fast_ mediated control of the SGNs was obtained in minutes by diffusing (and washing out) the compound into the cochlea. Interestingly, neural responses were observed in response to light pulses with energies between 0.5 and 1 µJ, which is the lower range of the optogenetic approaches applied to the adult gerbil cochlea (CatCh: 1-2 µJ, (Wrobel *et al*, 2018); f-Chrimson : ∼ 5 µJ, (Huet *et al*, 2021). Additionally, maximal oCAP responses were obtained in response to light pulse as short as 80 µs, which is substantially shorter than reported for optogenetic tools in the cochlea (Keppeler *et al*, 2018; Wrobel *et al*, 2018; Mager *et al*, 2018; Bali *et al*, 2021). Finally, TCP_fast_ enabled neural responses up to stimulation rate of 1.5 kHz. While this observation should be confirmed by single SGNs recording, it is interesting to note that this performance exceeds that obtained with the fastest opsins to date (Chronos: (Keppeler *et al*, 2018); vf-Chrimson: (Bali *et al*, 2021) while presenting lower activation threshold (TCP_fast_ ∼1 µJ, Chronos-ES/TS and vf-Chrimson-ES/TS ∼ 6 µJ).

TCP_fast_ provides similar or better performance than current optogenetic approaches in terms of speed and light requirements, while avoiding the gene therapy required to optogenetically modify SGNs. Yet, long-term availability, efficacy and safety of TCP_fast_ or more advanced photoswitches will need to be established. Moreover, neural degeneration often starts with the loss of postsynaptic structures (Kujawa & Liberman, 2009; Sergeyenko *et al*, 2013) limiting the utility of TCP_fast_ photopharmacology for optical cochlear implants. Future studies should focus on designing safe compound targeting channels expressed at the surface of the surviving SGN somas, while showing similar target specificity, fast kinetics and longer wavelength. This way, photopharmacology might establish a toolkit for photosensitization tailored to the specific neural status of the cochlea or other structures. In addition, a drug-based approach would appeal patients and the pharmaceutical industry and facilitate regulatory processes. Photopharmacology has previously demonstrated its potential in photosensitization of other sensory systems like the retina (Marc *et al*, 2014; Izquierdo-Serra *et al*, 2016; Tochitsky *et al*, 2017; Yue *et al*, 2016). To the best of our knowledge, this is the first proof-of-concept photopharmacology study in the auditory system and opens an avenue for auditory research and clinical applications such as in optical cochlear implants designed to fundamentally improve hearing restoration.

## ONLINE METHODS

### Click reaction

To a 1.5 ml glass vial containing azide **1** (‘head’, 1.00 mg, 1 eq) and copper(I) oxide (0.82 mg, 3 eq) in tetrahydrofuran (47 μl) and equipped with a magnetic stir bar was added a solution of ascorbic acid (1.34 mg, 4 eq) in water (94 μl) and the resulting mixture was vortexed for 1 min. Then, a solution of alkyne **2** (‘tail’, 0.47 mg, 1.1 eq) in tetrahydrofuran (47 μl) was added and the resulting mixture was stirred at room temperature for 45 min. The so-obtained final mixture was taken up in dimethylsulfoxide (193 μl), vortexed, centrifuged for 1 min to separate the insoluble copper(I) oxide particles, and finally divided into aliquots of the final compound stock solution (Supplementary Scheme S2).

Alternatively, the click reaction could be performed in an Eppendorf tube and stirred with a suitable mixer.

We observed that the catalytic performance of the copper(I) oxide may vary significantly from batch to batch, therefore the actual reaction time should be adjusted to obtain at least a 95% conversion of the starting material and a (TCP_fast_):(hydrolyzed TCP_fast_) ratio greater than 3 (Supplementary Figure S9). In vitro and in vivo biological characterizations were done using the same batch and results were reproducible across experiments.

### Rat hippocampal neural primary culture

All experiments were done in compliance with the national animal care guidelines and were approved by the board for animal welfare of the University Medical Center Goettingen and the animal welfare office of the state of Lower Saxony.

Primary hippocampal cultures were prepared from newborn P0-P3 pups Wistar rats. Brains were collected in a 10-cm petri dish containing ice-cold dissection media (HBSS (Gibco) + 10 mM Hepes (Gibco)). Hippocampi were separated from the brain, and meninges were removed. Hippocampi were digested with 2 ml pre-warmed 37°C 0.1% trypsin–EDTA (Gibco, Germany) for 20 min at 37°C. Trypsin was removed, and the tissue was washed three times with dissection medium. Dissection medium was replaced with 1 ml pre-warmed complete DMEM (DMEM with 1X Gluta-max, 10% FCS/FBS and 1% penicillin (100 U/ml)/streptomycin (100 µg/ml); all from Gibco). Tissue was triturated by gentle pipetting. The tissue suspension was filtered through a 100-µm cell strainer (BD Biosciences).

Cells were counted using the trypan blue exclusion method and cultured on 12-mm glass coverslips (Thermo Scientific) coated with poly-D-lysine (PDL, Sigma). Neurons were plated in 24-well plates at a density of 50,000 hippocampal neurons per coverslip in 500 µl of NB+ medium (Neurobasal with 2% B-27 supplement, 1% Glutamax and 1% penicillin (100 U/ml) /streptomycin (100 µg/ml); all from Gibco) at 37°C and 5% CO2. Half of the medium in each well was changed every 3-4 days.

### Electrophysiology recording conditions for rat hippocampal neurons

Before starting the recording, neurons between 10 to 18 days in vitro (div) were incubated with TCP_fast_ at concentrations: 25-50-75-100 µM (≤ 2% DMSO) for 2 min, in the absence of light and in pH 9 bath solution composed by (in mM): 100 NaCl, 1 MgCl_2_, 2.5 KCl, 2.5 CaCl_2_, 10 glucose and 50 sodiumcarbonate/sodiumbicarbonate, 310 mOsm/kg, pH 9 adjusted with NaOH. NMDG incubation solution at pH 9 is composed by (in mM): 100 NMDG, 2 MgCl_2_, 2.5 KCl, 10 glucose and 50 sodiumcarbonate/sodiumbicarbonate, 310 mOsm/kg, pH 9 adjusted with KOH.

Voltage and current-clamp recordings under whole-cell configuration were done using an Axopatch 200B amplifier, filtered at 5 kHz, digitized with an Axon DigiData 1440A interface (Axon Instruments). Acquisition software used was Clampex 10.5.2.6 (Axon Instruments).

Light stimulation was performed at saturating radiant flux (5-14 mW) using diode-pumped solid-state lasers (λ = 473 nm) focused into a 400-μm optic fiber. Light pulses were applied by a fast computer-controlled shutter (Uniblitz LS6ZM2, Vincent Associates, Rochester, USA). Radiant flux was adjusted by placement of density filters between the laser output and the optic fiber.

During recordings, neurons were maintained at room temperature (r.t., 25-27 °C) in a continuous perfusion of bath solution and clamped at -70 mV. Bath solution was composed of (in mM): 140 NaCl, 1 MgCl_2_, 2.5 KCl, 10 HEPES, 2.5 CaCl_2_ and 10–20 mM glucose to fix osmolarity to 310 mOsm/kg, pH 7.42 adjusted with NaOH. Borosilicate glass pipettes were pulled with a typical resistance of 3–8 MΩ for neurons. Pipette solution contained (in mM): 129 potassium gluconate, 10 HEPES, 10 KCl, 4 MgATP and 0.3 Na_3_GTP. Osmolarity is adjusted at 289 mOsm/kg and pH 7.2 adjusted with KOH.

### Drug preparation for in vivo cochlea infusion

Artificial perilymph solution consisted of the following (in mM): 137 NaCl; 5 KCl; 2 CaCl_2_; 1 MgCl_2_; 1 NaHCO_3_; 11 glucose; pH 7.4 adjusted with NaOH; osmolarity: 304 ± 4.3 mOsm/kg. Before each experiment, TCP_fast_ was diluted in artificial perilymph to a final concentration ranging from 2.5 to 12.5 μM.

### Animal preparation for cochlear potentials recordings and cochlear pharmacology

All experiments were done in compliance with the German national animal care guidelines and were approved by the board for animal welfare of the University Medical Center Göttingen and the animal welfare office of the state of Lower Saxony (agreement 2014/1726 and 2019/3188). Experiments were performed on adult (> 8 weeks old) Mongolian gerbils (*Meriones unguicalatus*) of both sexes.

Gerbils were anesthetized by isoflurane (5% for anesthesia induction, 1-2% for maintenance, frequent testing for the absence of hind-limb withdrawal reflex) and analgesia was obtained by subdermal injection of buprenorphine (0.1 mg/kg body weight) and carprofen (5 mg/kg body weight). Body temperature was maintained at 37 °C using a custom-designed heat plate. The cochlea was exposed following a retro-auricular approach and a bullostomy. The recording electrode was placed against the bony edge next to the round window (RW) leaving enough space to access the RW for pharmacological manipulation and optical stimulation. The pharmacological manipulation was made by filling the RW niche with the solution of interest. The RW membrane was punctured to increase the fluid exchange between the cochlea and RW niche. Between each solution, the RW niche was emptied by capillarity. The artificial perilymph solution had the following composition (in mM): 137 NaCl, 5 KCl, 2 CaCl2, 1 MgCl2, 1 NaHCO3, 11 glucose. The pH was ∼ 7.3 and osmolarity maintained at 300 mOsm/kg H2O.

### Acoustically evoked cochlear potentials

Acoustical cochlear potentials were obtained in response to 8 kHz tone burst (pulse duration = 8 ms, rise/fall time = 1 ms, repetition rate = 20 Hz, level: 20 to 80 dB SPL per 10 dB step, 200 repetitions per level). The mass potentials were amplified using a custom-made physiological amplifier and sampled at a rate of 50 kHz (NI PCI-6229, National Instrument). Stimulus generation and data acquisition were made using a custom-written software (MATLAB, MathWorks) employing National Instrument data acquisition cards in association with custom-build acoustic and laser-controller.

The cochlear microphonic was extracted by averaging the band-pass filtered (cut-off frequencies = 5.6 and 11.1 kHz) mass potential recorded using the RW electrode and its amplitude defined as the RMS value. The CAP and summating potentials were obtained by averaging the low-pass filtered (cut-off frequency = 3.5 kHz) mass-potential. The CAP amplitude was defined as the amplitude between the first negative peak (N_1_) and the following positive peak (P_1_). The summating potential amplitude was defined as the difference between the plateau response (between 5 and 7 ms) and the baseline prior to the stimulation onset.

### Optically evoked cochlear potentials

Optical cochlear potentials were obtained in response to blue light pulse delivered by a 200 µm optical fiber coupled to a 473 nm laser (MLL-FN-473-100, 100 mW, diode pumped solid state [DPSS]; Changchun New Industry Optoelectronics). Irradiance was calibrated with a laser power meter (LaserCheck; Coherent Inc.) The CAP was obtained was described above.

### Inner hair cell synapse counting

Cochleae were fixated in 4% formaldehyde for about 15 min, after which, they were decalcified in 0.12 M EDTA solution for about 12 hours (at 4°C). Organs of Corti were then: (i) isolated in phosphate buffer saline (PBS); (ii) incubated for an hour in Goat serum dilution buffer (GSDB, comprised of: 16% normal goat serum, 450 mM NaCl, 0.3% Triton X-100, 20 mM phosphate buffer, pH 7.4); (iii) treated for 2 hours with the primary antibodies: mouse anti-CtBP2 (BD Biosciences, 1:200), rabbit anti-myo6 (Proteus Biosciences, 1:200), chicken anti-Homer1 (SYsY, 1:200) for staining postsynapse, inner hair cells and presynapse, respectively. After washing the primary antibodies with PBS for 20 min, the secondary antibodies were incubated: Goat-anti-Chicken 488 (Invitrogen, 1:500), Goat-anti-Rabbit 568 (Thermo Fisher, 1:500), Goat-anti-Mouse 647 (Invitrogen, 1:500). After another washing step with PBS, samples were mounted on a glass slide in Mowiol (Carl Roth) mounting medium. Confocal images of the organ of Corti were first obtained using a 10x magnification (LSM 510 microscope, Carl Zeiss, Jena) in order to fit a Greenwood function (Greenwood, 1961; Müller, 1996) to them and localize the tonotopic position of the inner hair cells. The synapses were visualized at 0.5, 1, 2, 4, 8, 16 and 32 kHz using a 100x magnification (Abberior Instruments Expert Line STED microscope) and counted using the spot function from Imaris software (version 7.6.5).

### Data analysis and statistics

Amplitude of photocurrents were analyzed using IgorPro (Wavemetrics). Displayed whole-cell current traces have been filtered using the infinite impulse response digital filter from IgorPro (low-pass filter with cutoff of 50 Hz). The drift in current was corrected where appropriate with the IgorPro (WaveMetrics) software using a custom-made macro for drift correction. Statistics were done with OriginPro 8.5 (OriginLab) and Matlab (Mathworks). In vivo electrophysiological data were analyzed using custom-made Matlab routines.

## ACKNOWLEDGEMENTS

This research received funding from the European Union Research and Innovation Programme Horizon 2020 – Human Brain Project SG3 (945539), DEEPER (ICT-36-2020-101016787), Agency for Management of University and Research Grants/Generalitat de Catalunya (CERCA Programme; 2017-SGR-1442 project), Fonds Européen de Développement Économique et Régional (FEDER) funds, Ministry of Science and Innovation (Grant PID2019-111493RB-I00), Fundaluce and “la Caixa” foundations (ID 100010434, agreement LCF/PR/HR19/52160010). The project Clúster Emergent del Cervell Humà (CECH, 001-P-001682) is co-financed by the European Union Regional Development Fund within the framework of the ERDF Operational Program of Catalonia 2014-2020 with a grant of 50% of total eligible cost. A.G.-C. was supported by fellowship BES-2014–068169.

We thank Christiane Senger-Freitag, Sandra Gerke and Sina Langer for technical support, Gerhard Hoch for his expert technical support and Patricia Räke-Kügler for excellent administrative support. We thank Thomas Mager for providing the patch clamp setup. This work was funded by the European Research Council through the Advanced Grant “OptoHear” to TM under the European Union’s Horizon 2020 Research and Innovation program (grant agreement No. 670759), the Fraunhofer and Max-Planck Cooperation Program (NeurOpto grant) to TM, was further supported by the German Research Foundation through the Priority Program 1926 “Next generation optogenetics” to AH and TM, the Leibniz Program (to TM) and the Deutsche Forschungsgemeinschaft (DFG, German Research Foundation) under Germany’s Excellence Strategy -EXC 2067/1-390729940 to AH and TM. In addition, this research is supported by Fondation Pour l’Audition (FPA RD-2020-10).

We thank Dr. Vladimir Belov, Jurgen Bienert and Jan Seikowski for HPLC and LC-MS measurements of final TCP_fast_ products.

## AUTHOR CONTRIBUTIONS

AGC, AH, CM, AL, TM and PG designed the study. CM synthesised and analysed all compounds. AGC did electrophysiology in cultured neurons. AH did in vivo measurements in adult gerbils. AT did immunohistochemistry and image analysis. AL supervised the chemical synthesis. All authors contributed to the analysis of data and preparation of the article.

## COMPETING INTERESTS

TM is a co-founder and CEO of OptoGenTech company. The other authors declare no conflict of interests.

## 1. General methods and materials for chemical synthesis and characterization

All chemicals and solvents are from commercial suppliers and used without purification. All reactions were performed under inert atmosphere of argon. All analytical data of photoisomerizable compounds are given for the *trans* isomer unless otherwise stated. Reactions were monitored by thin layer chromatography (TLC: EMD/Millipore, silica gel 60 on aluminum support, layer thickness: 200 μm, particle size: 10-12 μm) by visualization under 254 and/or 365 nm lamp. Flash column chromatography: Panreac Silica Gel 60, 40-63 μm RE. NMR equipment and methods: Varian-Mercury 400 MHz & Varian VNMRS 500 MHz. Chemical shifts (δ) are reported in parts per million (ppm) against the reference compound tetramethylsilane using the signal of the residual non-deuterated solvent [Chloroform-*d* δ = 7.26 ppm (1H), δ = 77.16 ppm (13C); Dimethylsulfoxide-*d*_6_ δ = 2.50 ppm (1H), δ = 39.52 ppm (13C); Methanol-*d*_4_ δ = 3.31 ppm (1H), δ = 49.00 ppm (13C)]. HPLC-PDA-MS equipment and methods: Waters Alliance 2695 separation module coupled to a Waters 2996 photodiode detector (PDA) and a Waters ACQUITY QDa detector (single quadrupole; electrospray ionization), with the MassLynx software for data acquisition; SunFire C18 Column (100 Å, 5 μm, 4.6 mm X 150 mm); injection volume: 5 μL; mobile phase: water w/0.1% formic acid (solvent A) and acetonitrile w/0.1% formic acid (solvent B); elution method: flow 1 mL/min, gradient 0.0-1.0 min, 5% B; 1.0−7.0 min, 5−100% B; 7.0−8.0min, 100% B; 8.0−10.0 min, 100−5% B; runtime 10 min. Waters Alliance 2795 separation module coupled to a Waters 2996 photodiode detector (PDA) and a Waters 3100 Mass Detector (single quadrupole; electrospray ionization), with the MassLynx software for data acquisition; XSelect CSH C18 Column (130 Å, 3.5 µm, 4.6 mm X 50 mm); injection volume: 5 μL; mobile phase: water w/0.1% formic acid (solvent A) and acetonitrile w/0.1% formic acid (solvent B); elution method: flow 1.6 mL/min, gradient 0.0−3.5 min, 5−100% B; 3.5−4.5 min, 100% B; 4.5−5.0 min, 100−5% B; runtime 5 min. Spectra have been scanned between 200 and 800 Da with values every 0.1 seconds and peaks are given as mass/charge (*m*/*z*) ratio. Melting points of solid substances were determined on a Büchi melting point M-565 apparatus and are uncorrected. Optical rotations were measured with a Jasco P-2000 polarimeter operating on a sodium D-line (589 nm) at 25 °C, using a 10-cm path-length cell. High resolution mass spectrometry analyses were performed with a LTQ-FT Ultra Mass Spectrometer (Thermo Scientific) with NanoESI positive ionization. Each sample was reconstituted in MeOH and diluted with CH_3_CN/H_2_O/formic acid (50:50:1) for MS analysis. The sample was introduced by direct infusion (Automated Nanoelectrospray). The NanoMate (Advion BioSciences, Ithaca, NY, USA) aspirated the samples from a 384-well plate (protein Lobind) with disposable, conductive pipette tips, and infused the samples through the nanoESI Chip (which consists of 400 nozzles in a 20x20 array) towards the mass spectrometer. Spray voltage was 1.70 kV, delivery pressure 0.50 psi and *m*/*z* range 50-2000 Da. Data was acquired with Xcalibur software, vs.2.0SR2 (ThermoScientific). Elemental composition from experimental exact mass monoisotopic value was obtained with a dedicated algorithm integrated in Xcalibur software. Data are reported as mass-to-charge ratio (*m*/*z*) of the corresponding positively charged molecular ion.

## 2. Synthetic protocols for the preparation of the ‘head’ module

Compound **1** (‘head’ module) was prepared via a 5-step synthesis starting from commercially available materials (Scheme S1). Azobenzene **5** was obtained by reduction of its nitro precursor **4** with sodium sulfide nonahydrate. Pyroglutamate derivative **6**, prepared as previously described, (Volgraf *et al*, 2006) was coupled to compound **5** using HOBt/EDC activation to give the intermediate **7**, and then converted via mesylation into the corresponding azide derivative **8**. Hydrolysis of the pyroglutamate moiety with concomitant saponification of the ethyl ester provided the advanced intermediate **9**, which was finally converted into the desired compound **1** via removal of the *tert*-butoxycarbonyl protecting group under acidic conditions.

**Supplementary Scheme S1.**
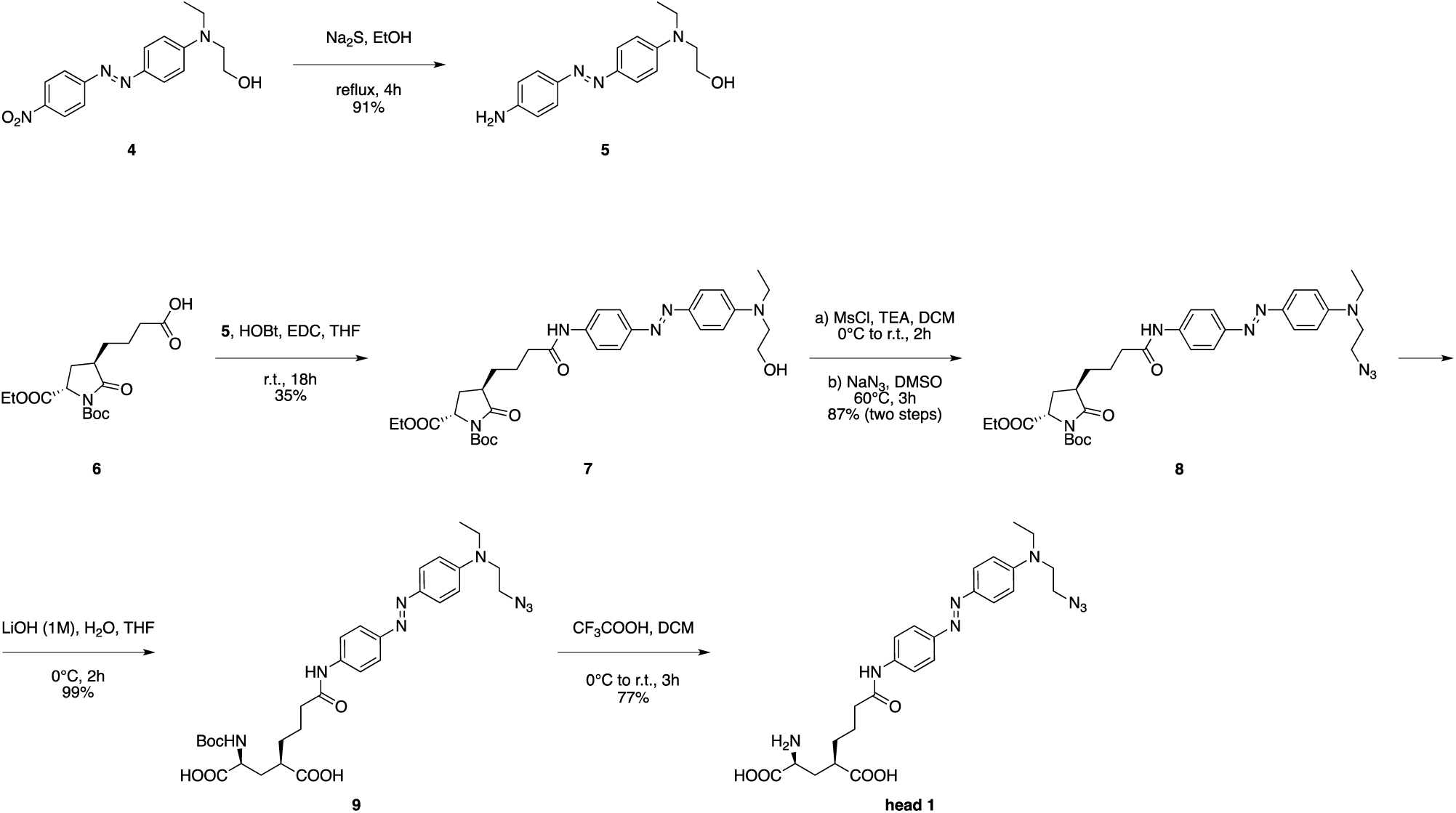
Chemical synthesis of the ‘head’ module (compound **1**)

**(*E*)-2-((4-((4-aminophenyl)diazenyl)phenyl)(ethyl)amino)ethanol [5]**

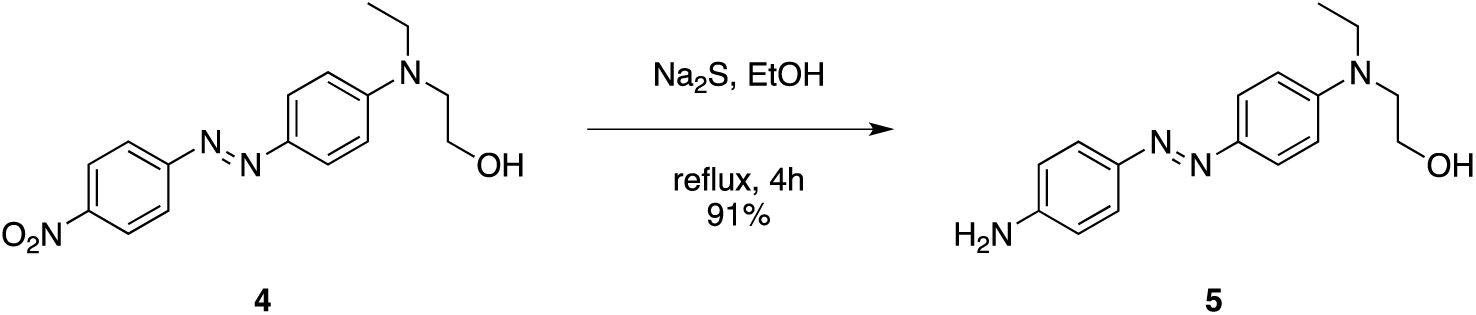

To a solution of (*E*)-2-(ethyl(4-((4-nitrophenyl)diazenyl)phenyl)amino)ethanol [**4**] (2.56 g, 8.14 mmol) in ethanol (250 mL) was added Na_2_S·9H_2_O (2.35 g, 9.77 mmol) and the reaction mixture was refluxed for 4 h. The mixture was then concentrated under reduced pressure and ethyl acetate (200 mL) was added to the residue. The organic layer was washed with water (200 mL) and brine (200 mL), dried over MgSO_4_, filtered and evaporated under reduced pressure to afford compound [**5**] as a dark red solid which was used in the next step without further purification (2.10 g, 91% yield).

*R*_f_ = 0.33 (TLC in cyclohexane/ethyl acetate = 1:1).

m.p. = 144 °C.

^1^H NMR (400 MHz, CD_3_OD) δ 7.73 – 7.66 (m, 2H), 7.64 – 7.58 (m, 2H), 6.83 – 6.77 (m, 2H), 6.77 – 6.71 (m, 2H), 3.74 (t, J = 6.4 Hz, 2H), 3.58 – 3.48 (m, 4H), 1.20 (t, J = 7.1 Hz, 3H).

^13^C NMR (101 MHz, CD_3_OD) δ 151.73, 150.98, 146.24, 144.62, 125.18, 124.95, 115.55, 112.46, 60.35, 53.43, 46.65, 12.45.

HRMS (*m*/*z*) calculated for C_16_H_21_N_4_O^+^ [M+H]^+^: 285.17099, found: 285.17032 (Δ_ppm_ = –2.34).

**(2*S*,4*R*)-1-*tert*-butyl 2-ethyl 4-(4-((4-((*E*)-(4-(ethyl(2-hydroxyethyl)amino)phenyl)diazenyl)phenyl) amino)-4-oxobutyl)-5-oxopyrrolidine-1,2-dicarboxylate [7]**

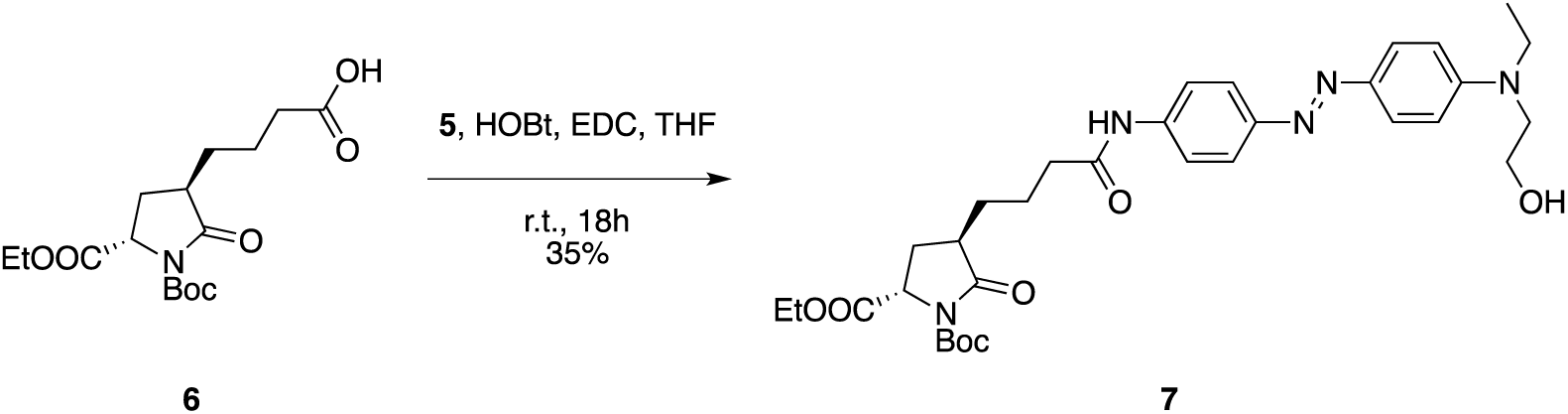

A suspension of 4-((3*R*,5*S*)-1-(*tert*-butoxycarbonyl)-5-(ethoxycarbonyl)-2-oxopyrrolidin-3-yl)butanoic acid [**6**] (250 mg, 0.73 mmol), 1-hydroxybenzotriazole hydrate (HOBt·xH_2_O, 223 mg, 1.46 mmol) and *N*-(3-dimethylaminopropyl)-*N*′-ethylcarbodiimide hydrochloride (EDC·HCl, 279 mg, 1.46 mmol) in anhydrous tetrahydrofuran (20 mL) was stirred at room temperature for 30 min. (*E*)-2-((4-((4-aminophenyl)diazenyl)phenyl)(ethyl)amino)ethanol [**5**] (311 mg, 1.09 mmol) was then added and the resulting mixture was stirred at room temperature for 18 h. The solvent was then evaporated and the residue was purified by direct flash chromatography (100% diethyl ether until complete elution of unreacted compound [**5**], then dichloromethane/methanol = 100:0 to 95:5 gradient) to afford compound [**7**] as a dark red oil (156 mg, 35% yield).

*R*_f_ = 0.34 (TLC in cyclohexane/ethyl acetate = 2:8).

[α]_D_ = –3.23 (c = 0.155, methanol).

^1^H NMR (400 MHz, CDCl_3_) δ 8.15 (s, 1H), 7.78 (dd, J = 8.9, 6.7 Hz, 4H), 7.64 (d, J = 8.8 Hz, 2H), 6.75 (d, J = 9.2 Hz, 2H), 4.55 (dd, J = 9.6, 1.5 Hz, 1H), 4.22 (q, J = 7.1 Hz, 2H), 3.83 (t, J = 6.0 Hz, 2H), 3.54 (t, J = 6.0 Hz, 2H), 3.48 (q, J = 7.0 Hz, 2H), 2.64 (dtd, J = 11.6, 8.5, 4.7 Hz, 1H), 2.51 (s, 1H), 2.39 (td, J = 7.2, 4.2 Hz, 2H), 2.24 (ddd, J = 13.2, 8.7, 1.5 Hz, 1H), 2.00 (ddd, J = 13.3, 11.7, 9.7 Hz, 1H), 1.94 – 1.82 (m, 1H), 1.76 (ddd, J = 13.7, 10.7, 6.9 Hz, 2H), 1.47 (s, 9H), 1.51 – 1.37 (m, 1H), 1.28 (t, J = 7.1 Hz, 3H), 1.19 (t, J = 7.0 Hz, 3H).

^13^C NMR (101 MHz, CDCl_3_) δ 175.75, 171.35, 171.26, 150.45, 149.51, 149.43, 143.67, 139.45, 125.10, 123.12, 120.08, 111.67, 83.87, 61.91, 60.21, 57.38, 52.51, 45.89, 41.64, 37.22, 29.76, 28.45, 28.00, 22.81, 14.32, 12.19.

HRMS (*m*/*z*) calculated for C_32_H_44_N_5_O_7_^+^ [M+H]^+^: 610.32353, found: 610.32276 (Δ_ppm_ = –1.25).

**(2*S*,4*R*)-1-*tert*-butyl 2-ethyl 4-(4-((4-((*E*)-(4-((2-azidoethyl)(ethyl)amino)phenyl)diazenyl)phenyl) amino)-4-oxobutyl)-5-oxopyrrolidine-1,2-dicarboxylate [8]**

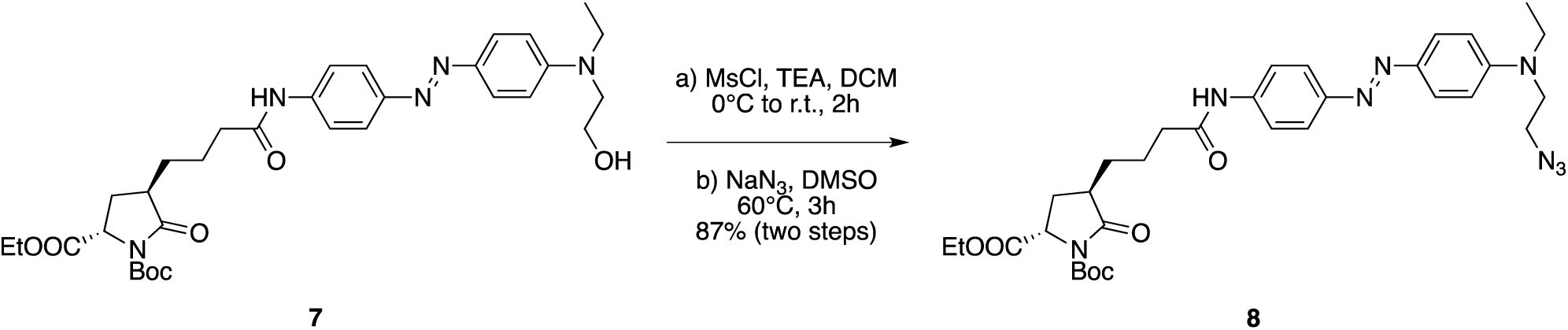

To a solution of (2*S*,4*R*)-1-*tert*-butyl 2-ethyl 4-(4-((4-((*E*)-(4-(ethyl(2-hydroxyethyl)amino)phenyl)diazenyl)phenyl)amino)-4-oxobutyl)-5-oxopyrrolidine-1,2-dicarboxylate [**7**] (136 mg, 0.22 mmol) in anhydrous dichloromethane (15 mL) at 0 °C was added triethylamine (187 μL, 1.34 mmol) followed by a slow addition of methanesulfonyl chloride (86 μL, 1.12 mmol). After 2 h of stirring at room temperature, the filtrate was concentrated to an oil and dissolved in dimethylsulfoxide (2 mL). Sodium azide (73 mg, 1.12 mmol) was then added and the mixture was stirred at 60 °C for 3 h in a sealed vessel. The reaction mixture was then allowed to cool to room temperature, diluted with brine (200 mL) and extracted with diethyl ether (3 × 200 mL). The combined organic layers were dried over MgSO_4_, filtered and evaporated to dryness. Purification by direct flash chromatography (*n*-hexane/ethyl acetate = 80:20 to 20:80 gradient) afforded compound [**8**] as an orange oil (123 mg, 87% yield).

*R*_f_ = 0.58 (TLC in cyclohexane/ethyl acetate = 2:8).

[α]_D_ = –2.63 (c = 0.165, methanol).

^1^H NMR (400 MHz, CDCl_3_) δ 7.94 (s, 1H), 7.83 (dd, J = 9.9, 9.0 Hz, 4H), 7.68 (d, J = 8.8 Hz, 2H), 6.75 (d, J = 9.2 Hz, 2H), 4.57 (dd, J = 9.6, 1.4 Hz, 1H), 4.23 (q, J = 7.1 Hz, 2H), 3.58 (t, J = 6.0 Hz, 2H), 3.55 – 3.47 (m, 4H), 2.67 (dtd, J = 11.7, 8.4, 4.8 Hz, 1H), 2.43 (td, J = 7.2, 5.0 Hz, 2H), 2.27 (ddd, J = 13.3, 8.7, 1.5 Hz, 1H), 2.02 (ddd, J = 13.3, 11.7, 9.6 Hz, 1H), 1.93 (dtd, J = 17.7, 7.4, 4.8 Hz, 1H), 1.87 – 1.74 (m, 2H), 1.58 – 1.44 (m, 1H), 1.49 (s, 9H), 1.29 (t, J = 7.1 Hz, 3H), 1.23 (t, J = 7.1 Hz, 3H).

^13^C NMR (101 MHz, CDCl_3_) δ 175.57, 171.36, 171.03, 149.54, 149.49, 149.47, 144.06, 139.54, 125.14, 123.26, 119.98, 111.63, 83.81, 61.90, 57.35, 49.70, 49.10, 45.92, 41.67, 37.31, 29.78, 28.53, 28.03, 22.84, 14.34, 12.42.

HRMS (*m*/*z*) calculated for C_32_H_43_N_8_O_6_^+^ [M+H]^+^: 635.33001, found: 635.32940 (Δ_ppm_ = –0.96).

**(2*R*,4*S*)-2-(4-((4-((*E*)-(4-((2-azidoethyl)(ethyl)amino)phenyl)diazenyl)phenyl)amino)-4-oxobutyl)-4-((*tert*-butoxycarbonyl)amino)pentanedioic acid [9]**

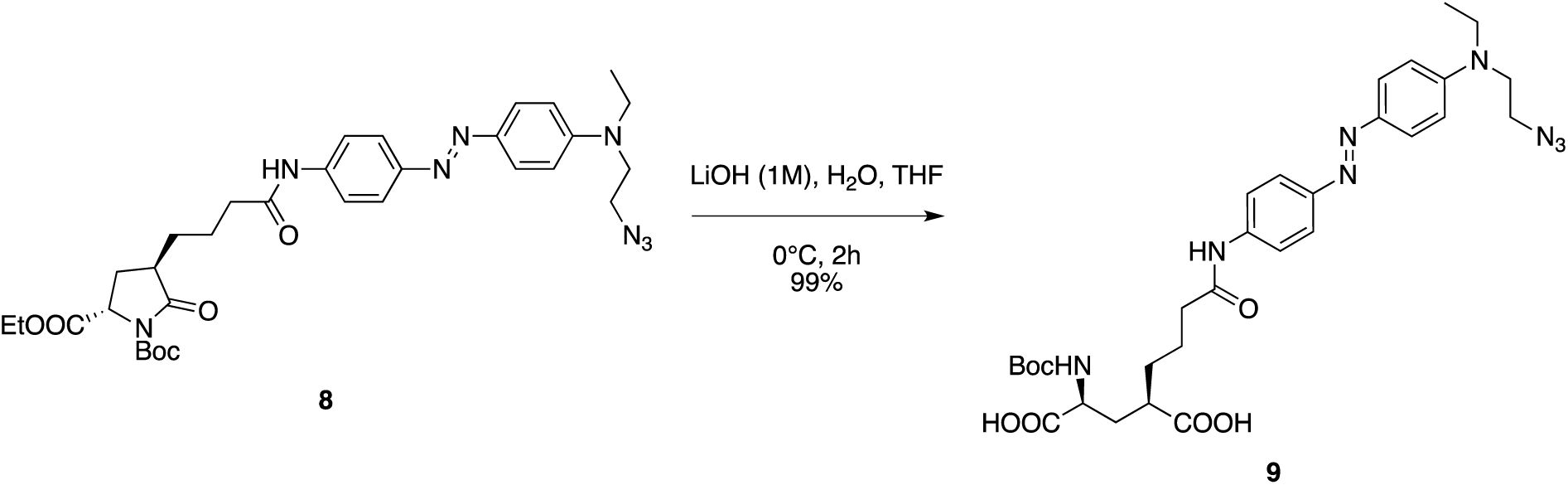

To a solution of (2*S*,4*R*)-1-*tert*-butyl 2-ethyl 4-(4-((4-((*E*)-(4-((2-azidoethyl)(ethyl)amino)phenyl)diazenyl)phenyl)amino)-4-oxobutyl)-5-oxopyrrolidine-1,2-dicarboxylate [**8**] (133 mg, 0.21 mmol) in tetrahydrofuran (6.3 mL) at 0 °C was added a 1.0 M aqueous solution of lithium hydroxide (6.3 mL) and the resulting mixture was stirred at 0 °C for 2 h. The reaction was then acidified to pH 2 with 1.0 M H_2_SO_4_ and extracted with ethyl acetate (5 × 30 mL). The combined organic layers were dried over MgSO_4_, filtered and evaporated to dryness to afford compound [**9**] as a dark red oil which was used in the next step without further purification (130 mg, 99% yield).

*R*_f_ = 0.55 (TLC in dichloromethane/methanol = 8:2).

[α]_D_ = +14.57 (c = 0.270, methanol).

^1^H NMR (400 MHz, CD_3_OD) δ 7.81 (d, J = 9.2 Hz, 2H), 7.78 (d, J = 8.9 Hz, 2H), 7.70 (d, J = 9.0 Hz, 2H), 6.87 (d, J = 9.3 Hz, 2H), 4.17 – 4.06 (m, 1H), 3.64 (t, J = 6.1 Hz, 2H), 3.61 – 3.50 (m, 4H), 2.65 – 2.50 (m, 1H), 2.43 (t, J = 7.1 Hz, 2H), 2.30 – 2.12 (m, 1H), 2.10 – 1.89 (m, 1H), 1.88 – 1.67 (m, 4H), 1.67 – 1.55 (m, 1H), 1.44 (s, 9H), 1.24 (t, J = 7.0 Hz, 3H).

^13^C NMR (101 MHz, CD_3_OD) δ 178.71, 176.05, 174.15, 151.36, 150.63, 144.94, 141.34, 125.98, 123.80, 121.22, 112.74, 80.49, 53.56, 50.47, 50.27, 46.49, 43.15, 37.74, 34.90, 33.44, 30.13, 28.73, 24.46, 12.46.

HRMS (*m*/*z*) calculated for C_30_H_41_N_8_O_7_^+^ [M+H]^+^: 625.30927, found: 625.30847 (Δ_ppm_ = –1.28).

**(2*S*,4*R*)-2-amino-4-(4-((4-((*E*)-(4-((2-azidoethyl)(ethyl)amino)phenyl)diazenyl)phenyl)amino)-4-oxobutyl)pentanedioic acid [1]**

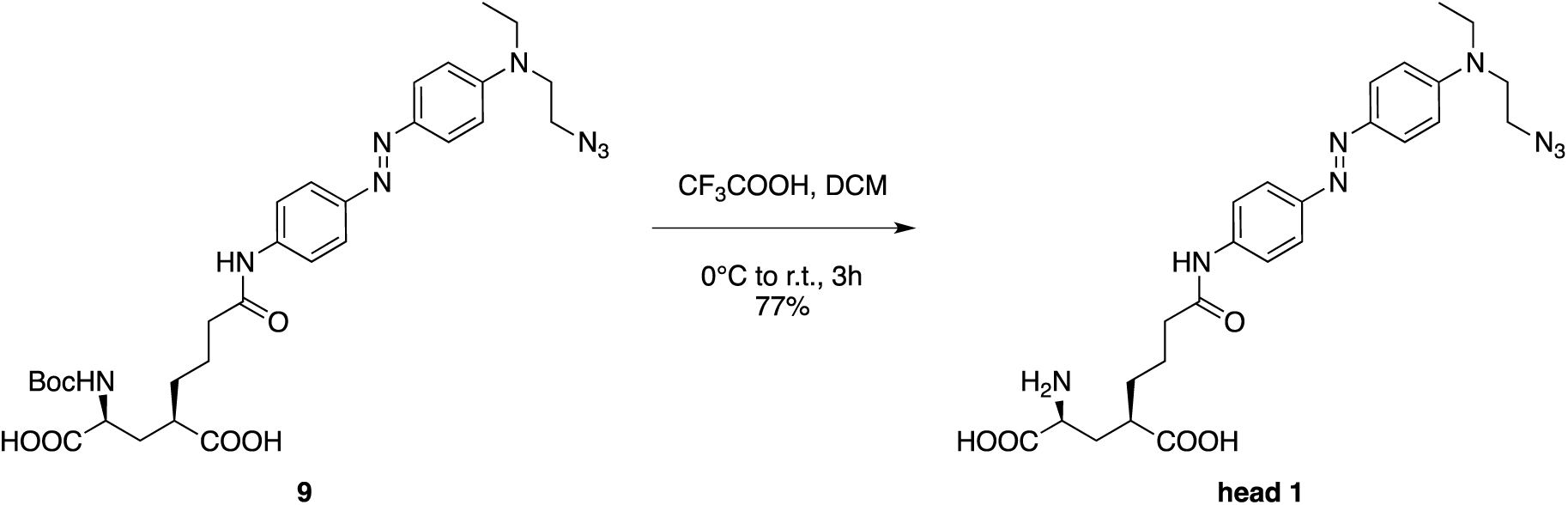

Trifluoroacetic acid (313 μL, 4.09 mmol) was added dropwise to a solution of (2*R*,4*S*)-2-(4-((4-((*E*)-(4-((2-azidoethyl)(ethyl)amino)phenyl)diazenyl)phenyl)amino)-4-oxobutyl)-4-((*tert*-butoxycarbonyl)amino)pentanedioic acid [**9**] (128 mg, 0.20 mmol) in anhydrous DCM (15 mL) at 0 °C, and the resulting solution was stirred at room temperature for 3 h. The solvent was then removed under reduced pressure and the residue was purified by reversed-phase flash chromatography (water/acetonitrile = 100:0 to 0:100 gradient) to afford compound [**1**] as an orange solid (83 mg, 77% yield; purity ≥99% as determined by HPLC-PDA analysis).

*R*_f_ = 0.10 (TLC in dichloromethane/methanol = 8:2).

m.p. = 150 °C (dec.).

[α]_D_ = +11.70 (c = 0.265, methanol).

^1^H NMR (500 MHz, CD_3_OD) δ 7.84 – 7.78 (m, 2H), 7.81 – 7.74 (m, 2H), 7.74 – 7.68 (m, 2H), 6.91 – 6.83 (m, 2H), 3.69 – 3.61 (m, 3H), 3.61 – 3.51 (m, 4H), 2.68 – 2.59 (m, 1H), 2.48 – 2.41 (m, 2H), 2.28 (ddd, J = 14.7, 9.0, 4.7 Hz, 1H), 1.93 (ddd, J = 14.7, 8.5, 4.7 Hz, 1H), 1.86 – 1.72 (m, 4H), 1.72 – 1.61 (m, 2H), 1.24 (t, J = 7.0 Hz, 3H).

^13^C NMR (126 MHz, CD_3_OD) δ 179.72, 174.28, 173.95, 151.35, 150.61, 144.94, 141.39, 125.97, 123.79, 121.21, 112.74, 54.39, 50.48, 50.28, 46.50, 43.51, 37.74, 33.96, 32.71, 24.40, 12.46.

UV-Vis (PBS): λ_max_ (abs) = 460 nm, ε = 2.927 × 10^4^ M^−1^·cm^−1^.

*R*_t_ (HPLC-PDA, XSelect CSH C18 Column) = 2.25 min (*trans* isomer).

HRMS (*m*/*z*) calculated for C_25_H_33_N_8_O_5_^+^ [M+H]^+^: 525.25684, found: 525.25883 (Δ_ppm_ = +3.78).

## 3. Supplementary Figures

**Supplementary Figure S1.**
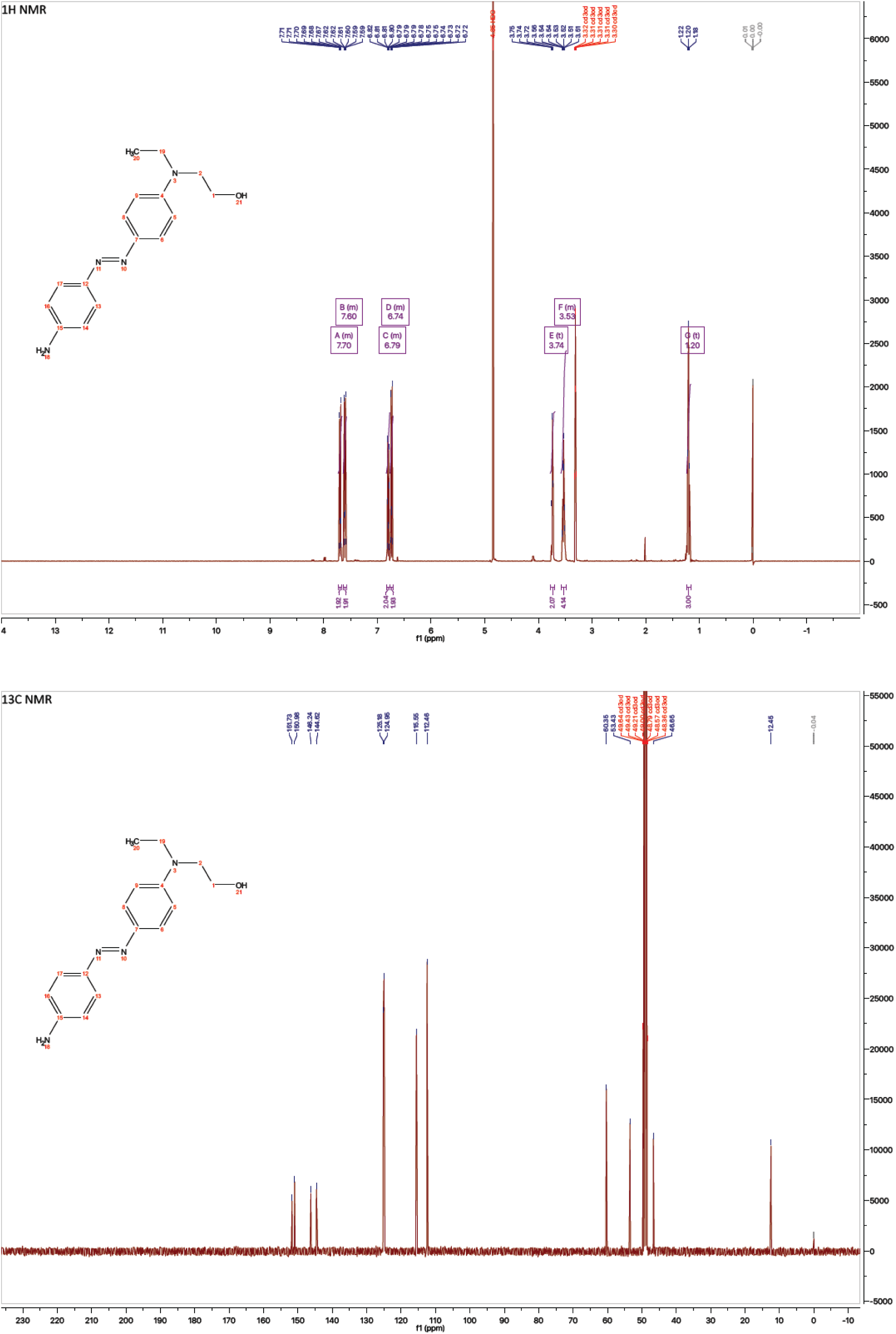
NMR spectra of compound 5.

**Supplementary Figure S2.**
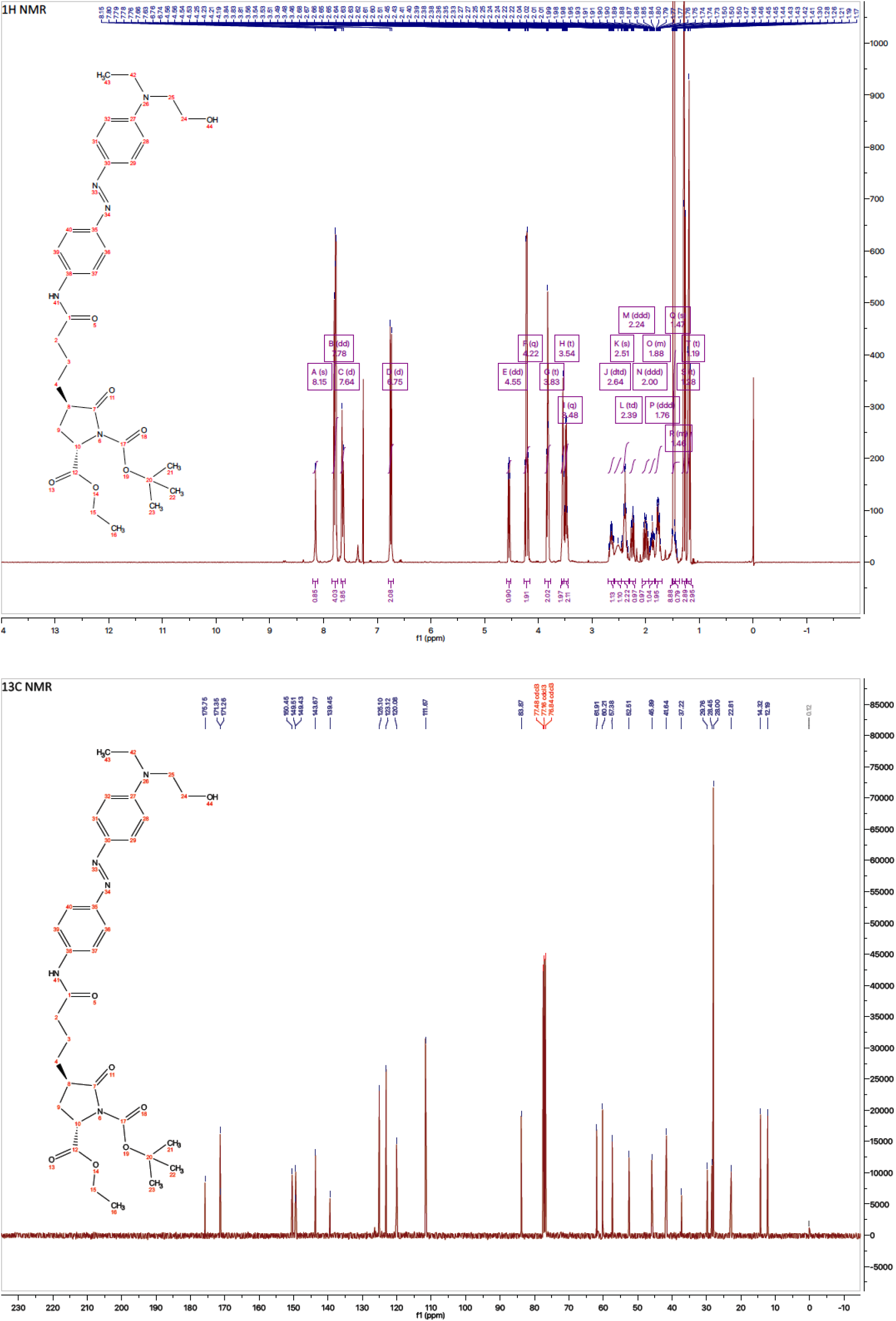
NMR spectra of compound 7.

**Supplementary Figure S3.**
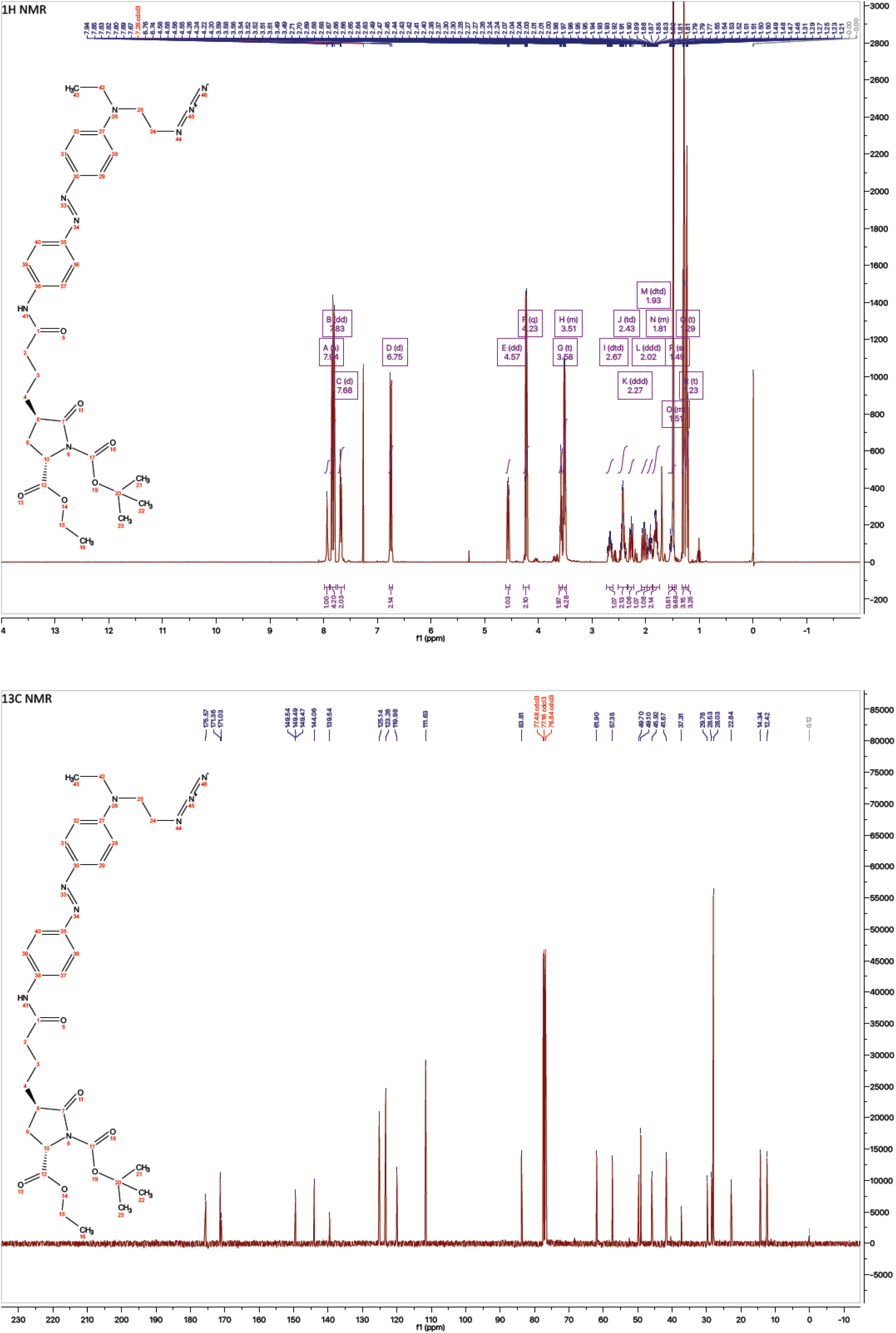
NMR spectra of compound 8.

**Supplementary Figure S4.**
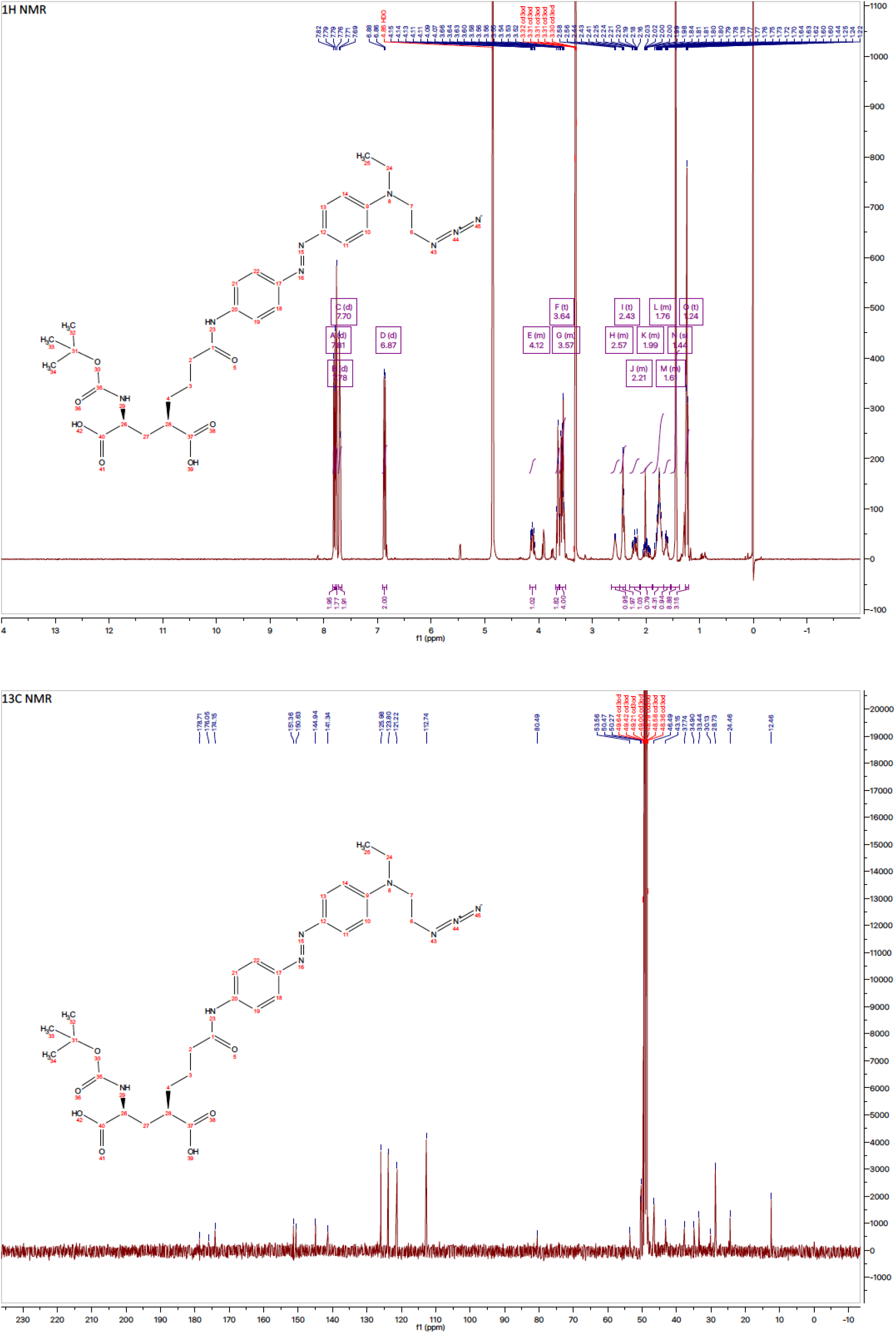
NMR spectra of compound 9.

**Supplementary Figure S5.**
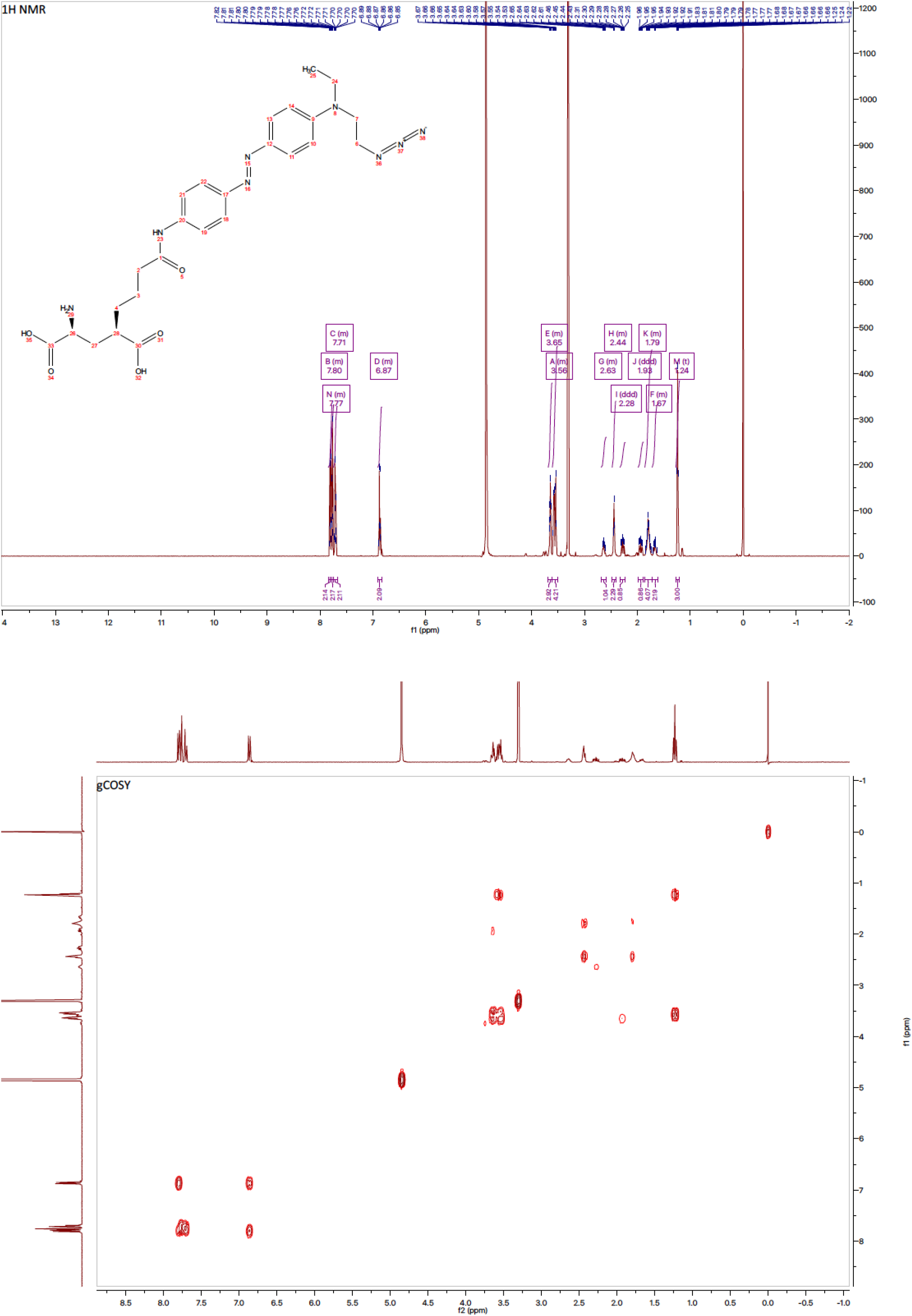
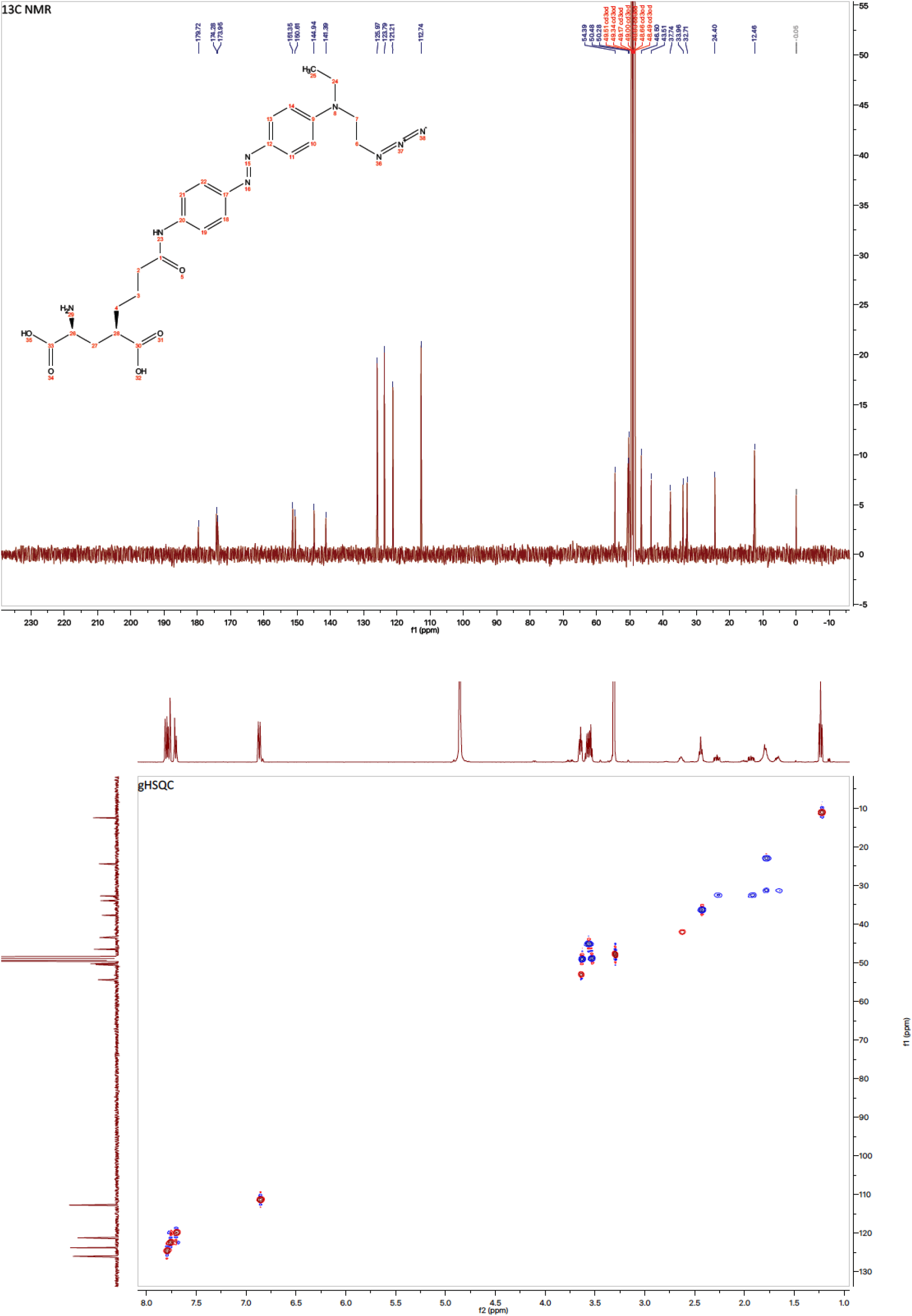
NMR spectra of compound 1.

**Supplementary Figure S6.**
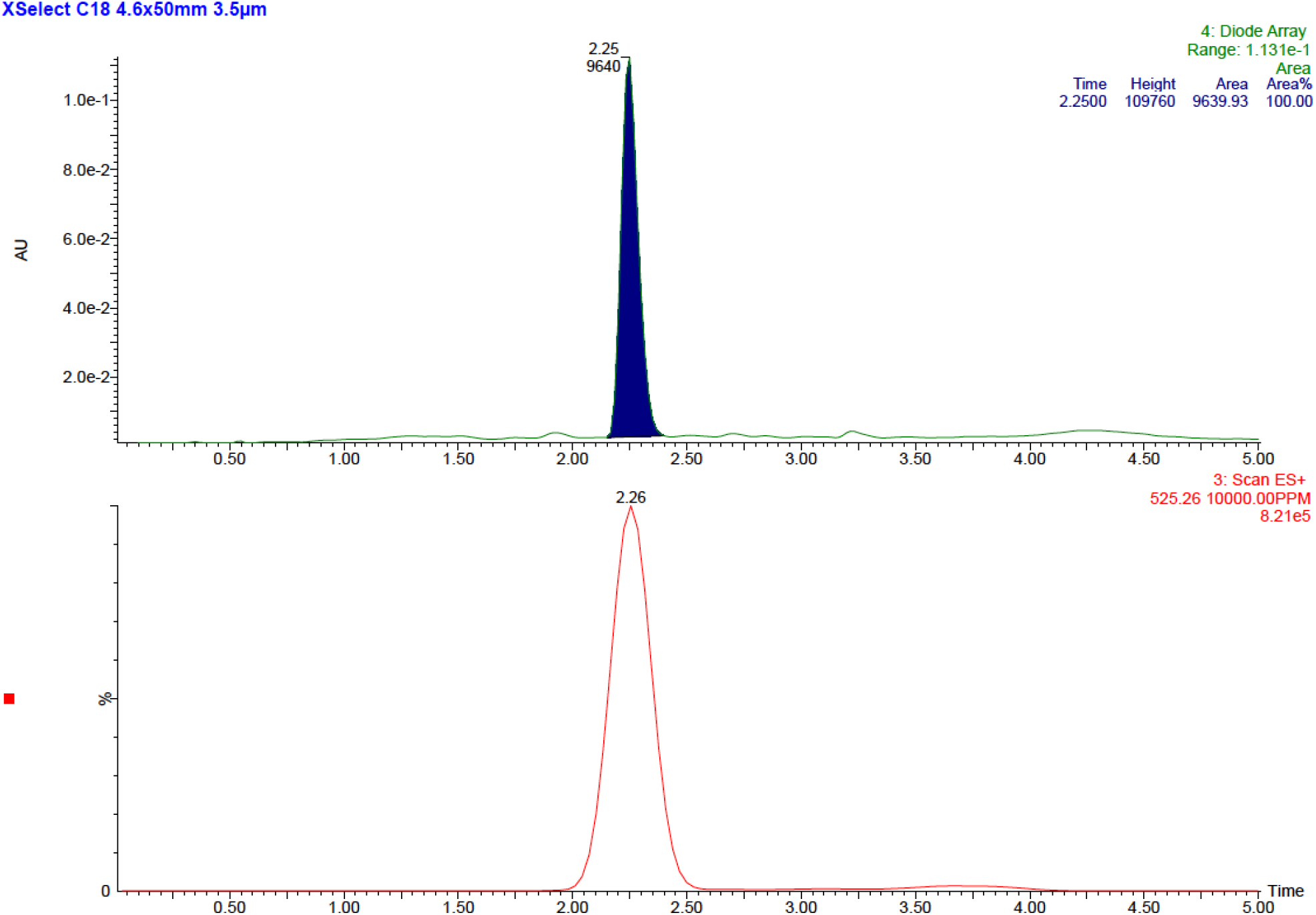
HPLC chromatogram of compound 1.

**Supplementary Figure S7.**
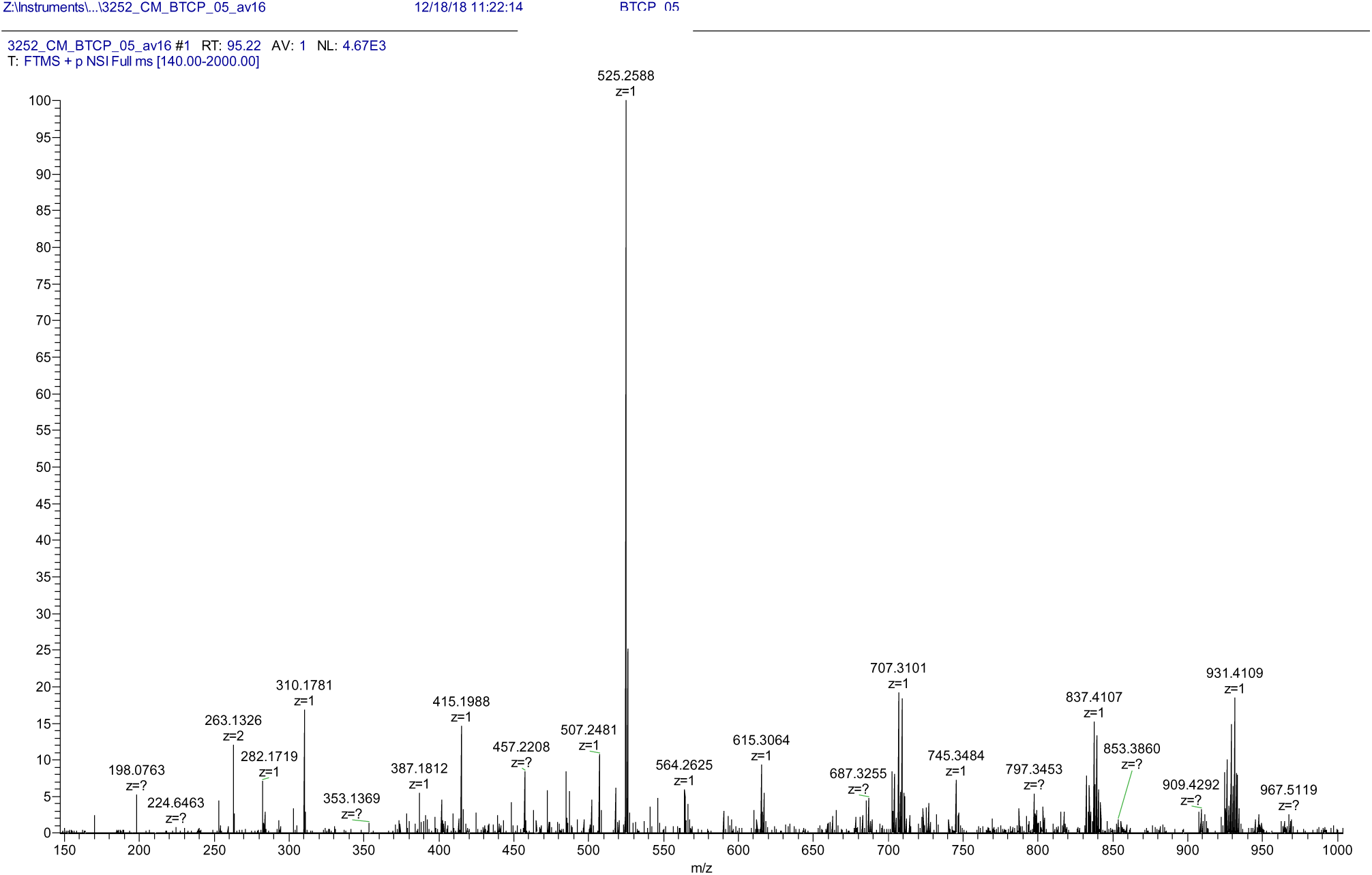
High-resolution mass spectrum of compound 1.

**Supplementary Figure S8.**
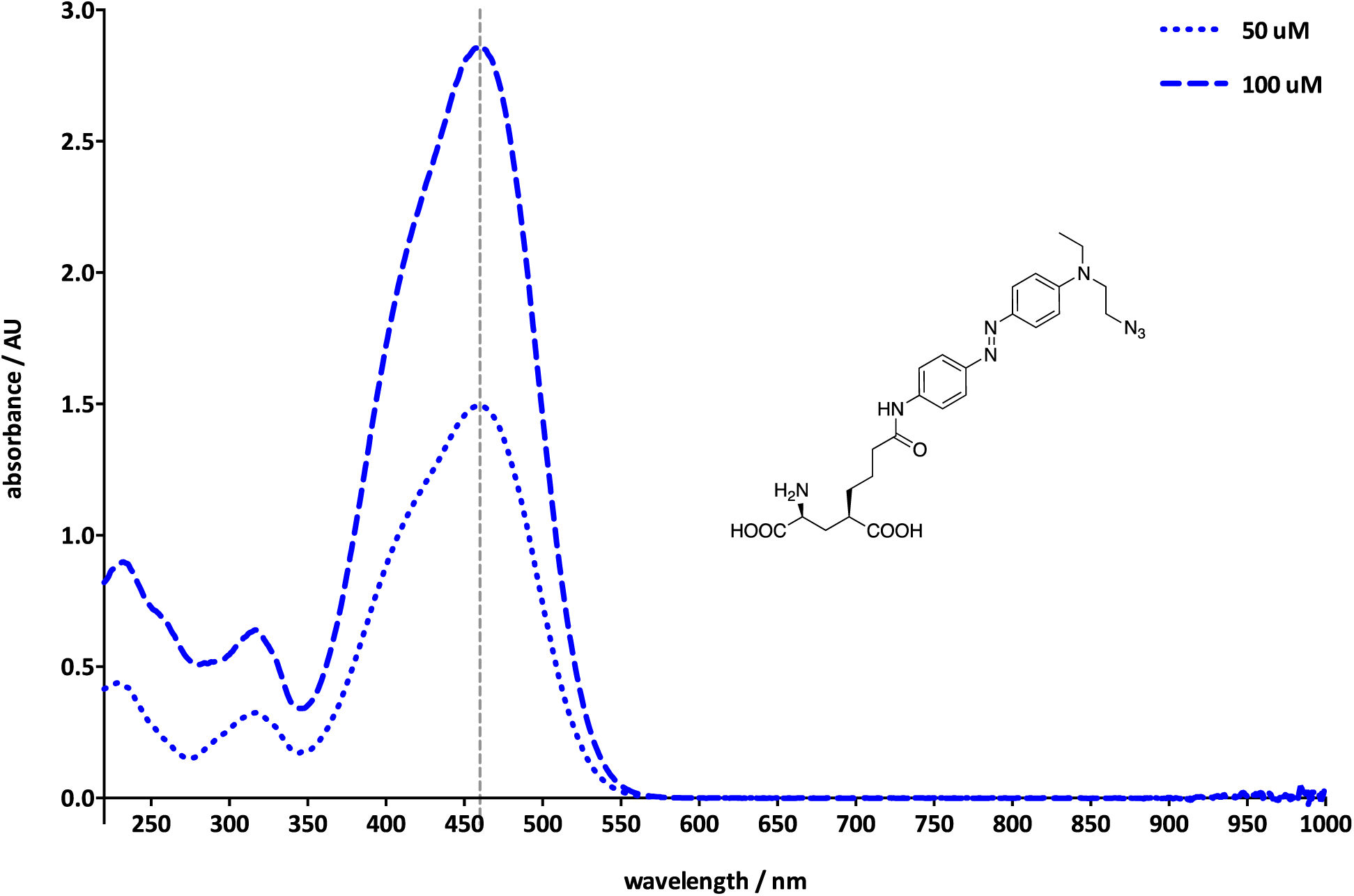
UV-Vis absorption spectrum of compound 1 (*trans* isomer) in PBS buffer at pH 7.4.

## 4. Synthetic protocol for the preparation of TCPfast

Head (**1**) and tail (**2**, commercially available) were combined to form the final TCP_fast_ (**3**) compound using a “click” version of the Huisgen azide-alkyne 1,3-dipolar cycloaddition.(Himo *et al*, 2005) Since such NHS-ester derivatives are very short-lived, we characterized the products of the click reaction crude of TCP_fast_ after subsequent reaction with pure L-lysine as a mock protein residue. Detailed analysis by LC-MS confirmed the presence of the intended TCP_fast_–lysine adduct with an intact glutamate moiety and of some of the expected byproducts.

### General procedure

To a 1.5 ml glass vial containing azide **1** (‘head’, 1.00 mg, 1 eq) and copper(I) oxide (0.82 mg, 3 eq) in tetrahydrofuran (47 μL) and equipped with a magnetic stir bar was added a solution of ascorbic acid (1.34 mg, 4 eq) in water (94 μL) and the resulting mixture was vortexed for 1 min. Then, a solution of alkyne **2** (‘tail’, 0.47 mg, 1.1 eq) in tetrahydrofuran (47 μL) was added and the resulting mixture was stirred at room temperature for 45 min. The so-obtained final mixture was taken up in dimethylsulfoxide (193 μL), vortexed, centrifuged for 1 min to separate the insoluble copper(I) oxide particles, and finally divided into aliquots of the final compound stock solution (Supplementary Scheme S2).

We observed that the catalytic performance of the copper(I) oxide may vary significantly from batch to batch, therefore the actual reaction time should be adjusted accordingly in order to obtain a ≥95% conversion of the starting material and a (TCP_fast_):(hydrolyzed TCP_fast_) ratio greater than 3 (Supplementary Figure S9).

**Supplementary Scheme S2.**
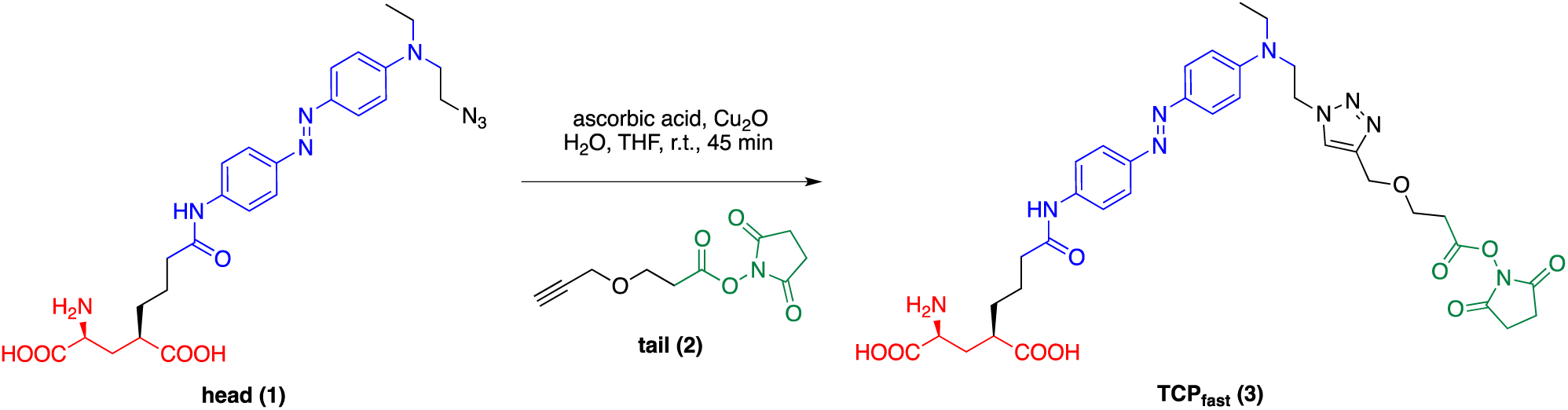
Preparation of TCP_fast_ via Cu(I)-catalyzed azide-alkyne 1,3-dipolar cycloaddition (“click”) reaction.

**Supplementary Figure S9.**
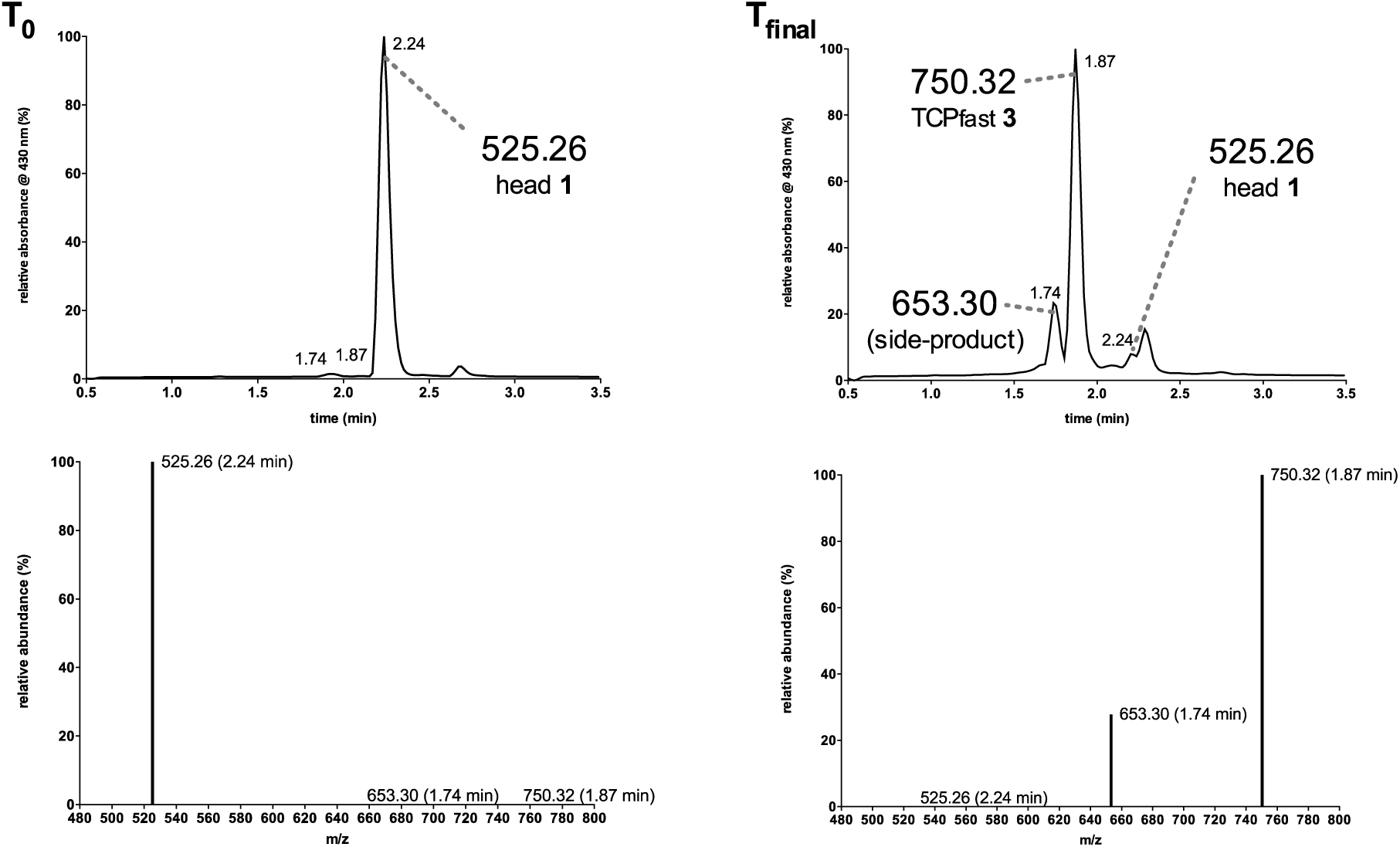
Analysis of the initial and final mixture of a representative “click” reaction for the preparation of TCP_fast_ (**3**), showing the conversion of the starting material to the final desired product and its side-product [top: HPLC chromatograms; *y*-axis shows relative absorbance (%) at 430 nm; bottom: selected ion recording (SIR) of the corresponding HPLC chromatograms for head (**1**), TCP_fast_ (**3**) and its major side-product; *y*-axis shows relative abundance (%) of the ion species; see also Supplementary Scheme S3 for further details].

**Supplementary Figure S10.**
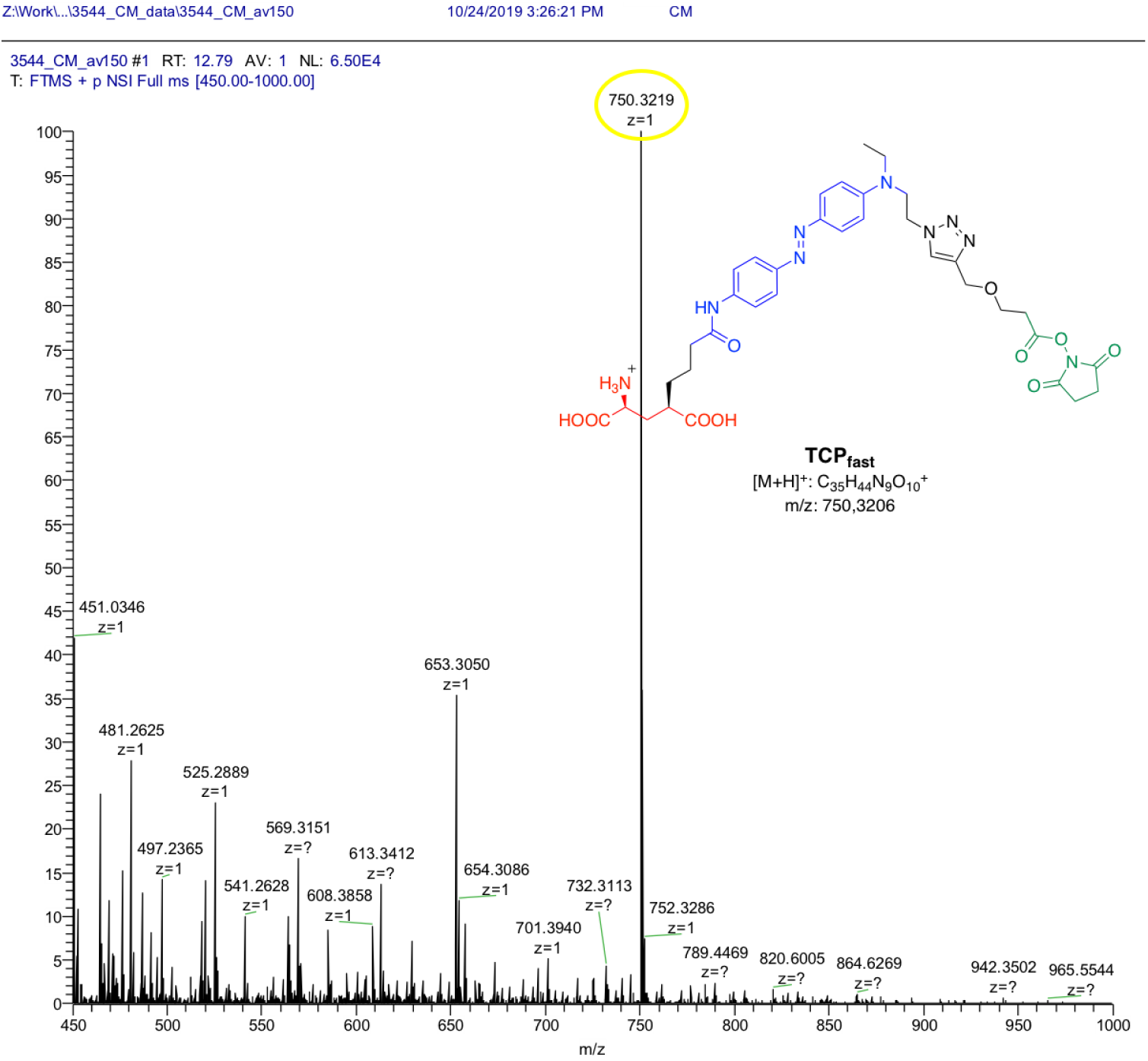
High-resolution mass spectrum of TCP_fast_ (**3**).

## 5. Characterization of TCPfast, lysine-adduct, and side-product by LC-MS

**Supplementary Scheme S3.**
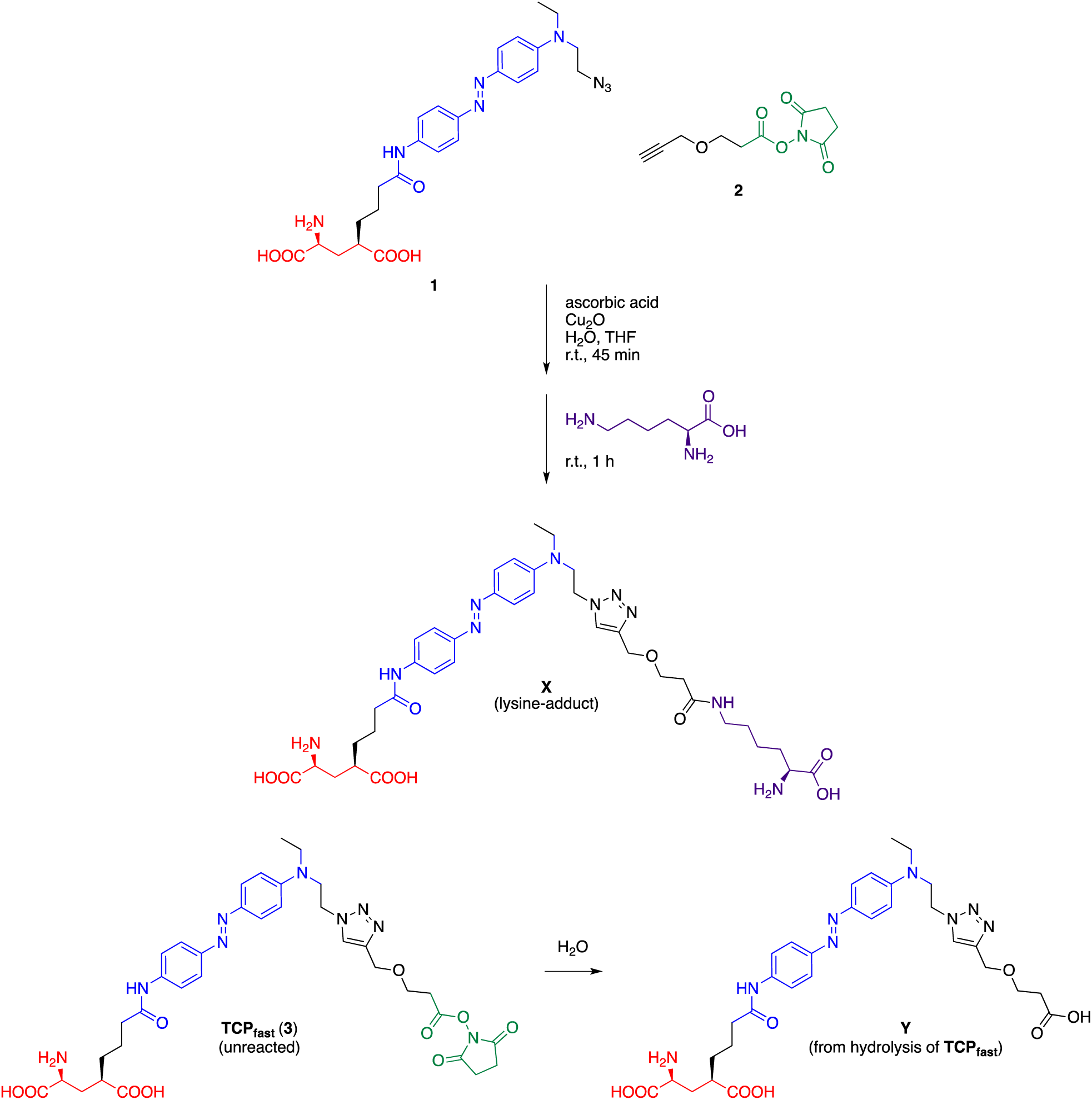
Generation and structure of the TCP_fast_–lysine adduct (**X**) and detected main side-product (**Y**).

**Supplementary Table S1.** Calculated mass-to-charge (*m*/*z*) ratio for compounds **1**, TCP_fast_ (**3**), **X**, **Y**.

| Compound | $m/z$<br>calculated<br>for $[M+H]^+$ | $m/z$<br>calculated for<br>$[M+2H]^{2+}$ |
| --- | --- | --- |
| **1** | $C_{25}H_{33}N_8O_5^+$:<br>525.26 | $C_{25}H_{34}N_8O_5^{2+}$:<br>263.13 |
| **TCP<sub>fast</sub> (3)** | $C_{35}H_{44}N_9O_{10}^+$:<br>750.32 | $C_{35}H_{45}N_9O_{10}^{2+}$:<br>375.66 |
| **X** | $C_{37}H_{53}N_{10}O_9^+$:<br>781.40 | $C_{37}H_{54}N_{10}O_9^{2+}$:<br>391.20 |
| **Y** | $C_{31}H_{41}N_8O_8^+$:<br>653.30 | $C_{31}H_{42}N_8O_8^{2+}$:<br>327.16 |

**Supplementary Figure S11.**
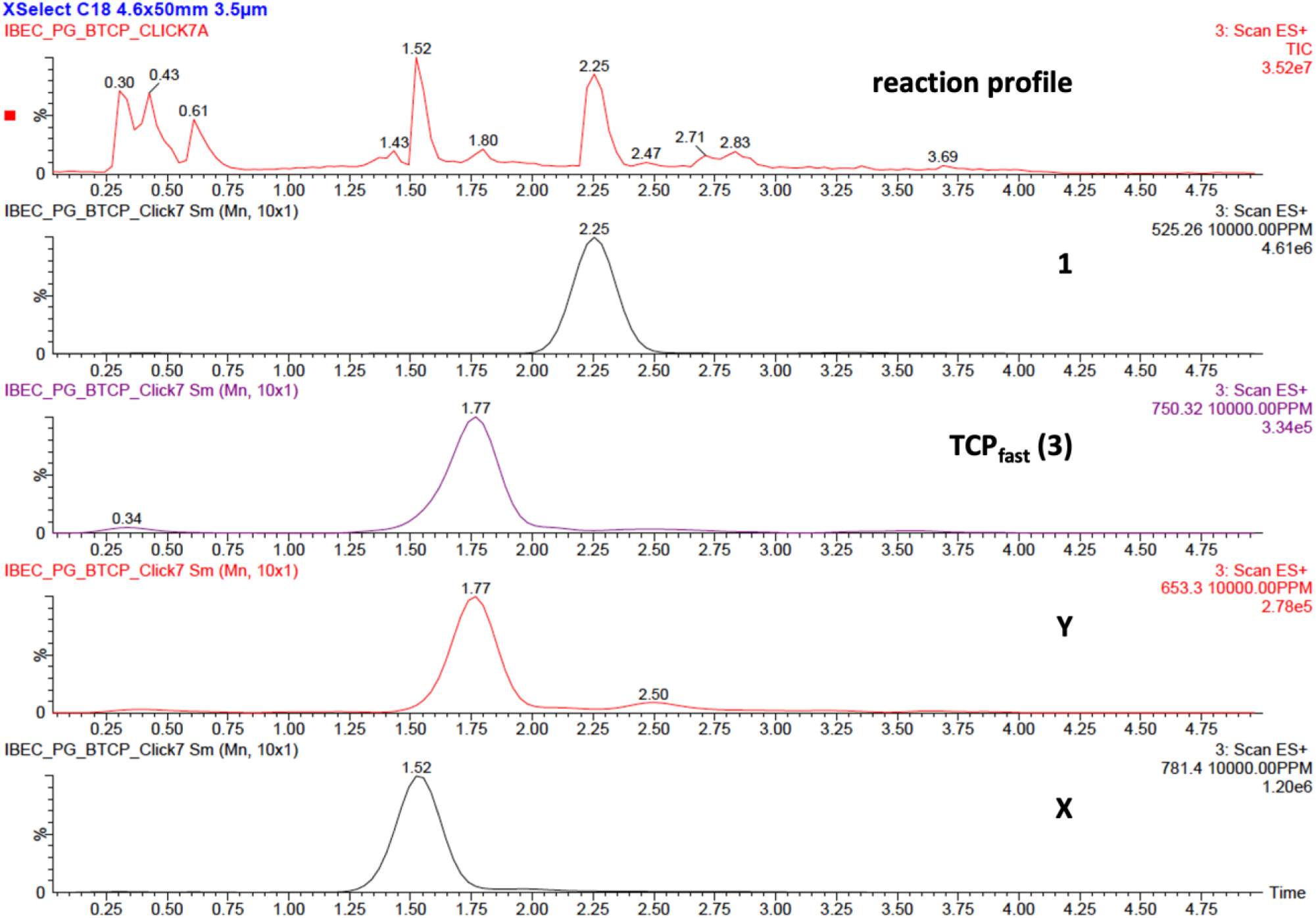
LC-MS analysis of the final mixture for the reaction illustrated in Scheme S3 (arbitrary case).

**Supplementary Figure S12.**
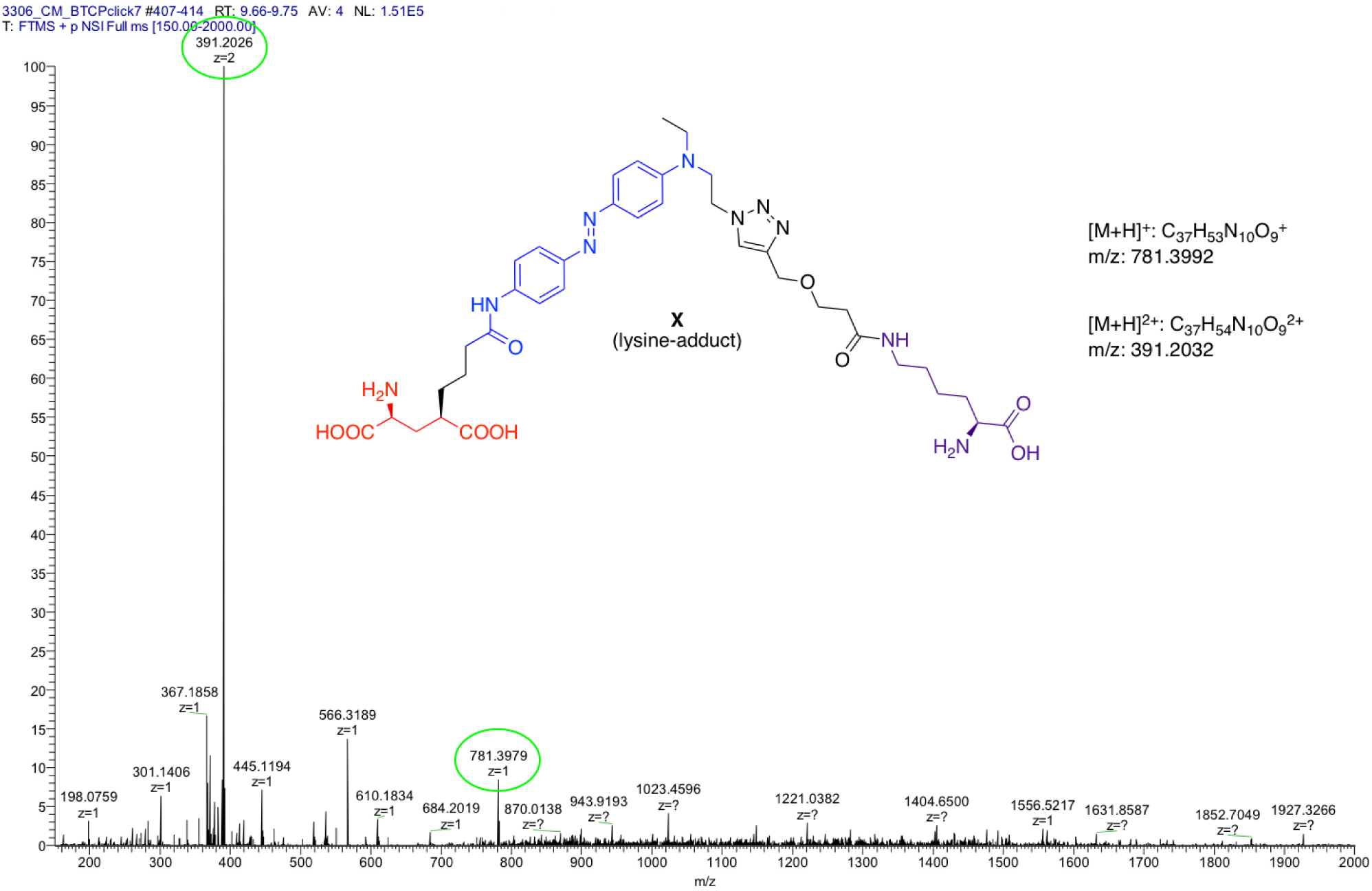
High-resolution mass spectrum of the TCP_fast_–lysine adduct (**X**).

## 6. Characterization of TCPfast in cultured neurons

**Supplementary Figure S13.**
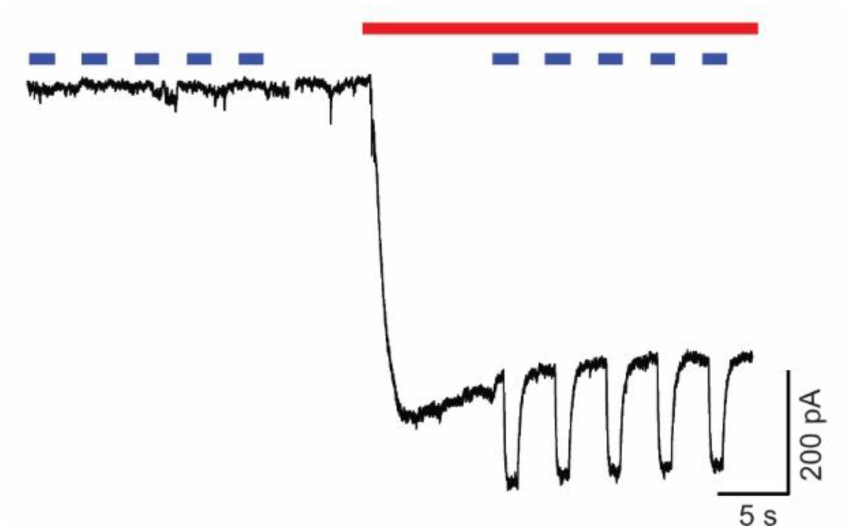
TCP_fast_ conjugation produces photocurrents in untransfected hippocampal neurons in the presence of glutamate. Current recording in whole cell voltage clamp mode in dissociated hippocampal neurons after incubation in TCP_fast_ (100 μM for 2 min at pH9). Inward current induced by bath perfusion of 300 µM glutamate (red bar) and 473 nm light (blue bars, 1 s). Time gap between traces corresponds to immediate subsequent recordings (< 2s).

**Supplementary Figure S14.**
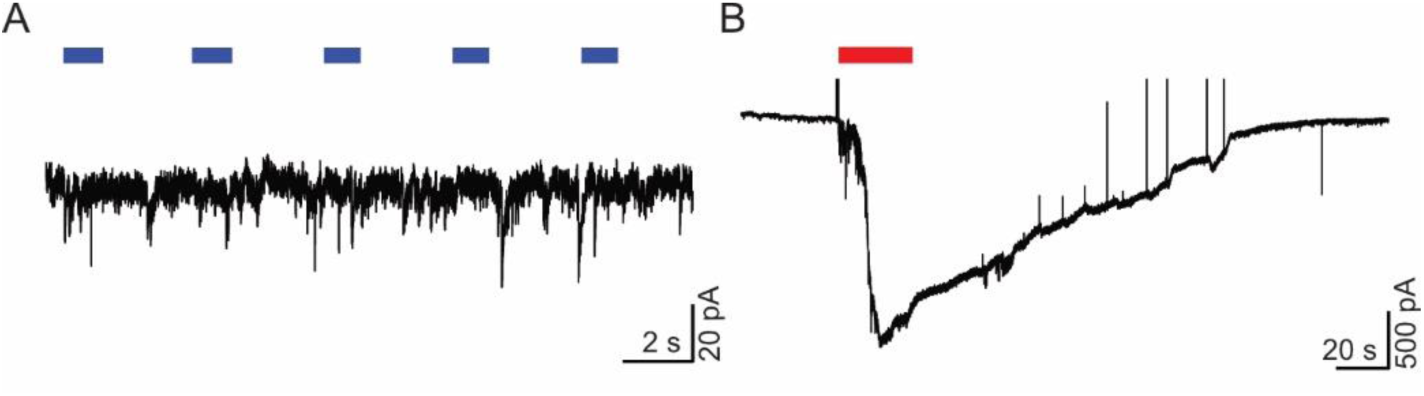
Light stimulation does not affect neuronal responses prior to TCP_fast_ incubation. **A.** Hippocampal neurons non-incubated with TCP_fast_ do not respond to light (473 nm for 1 s, blue bars). **B.** Physiological responses to glutamate perfusion (300 µM, red bar) indicate expression of glutamatergic receptors.

**Supplementary Figure S15.**
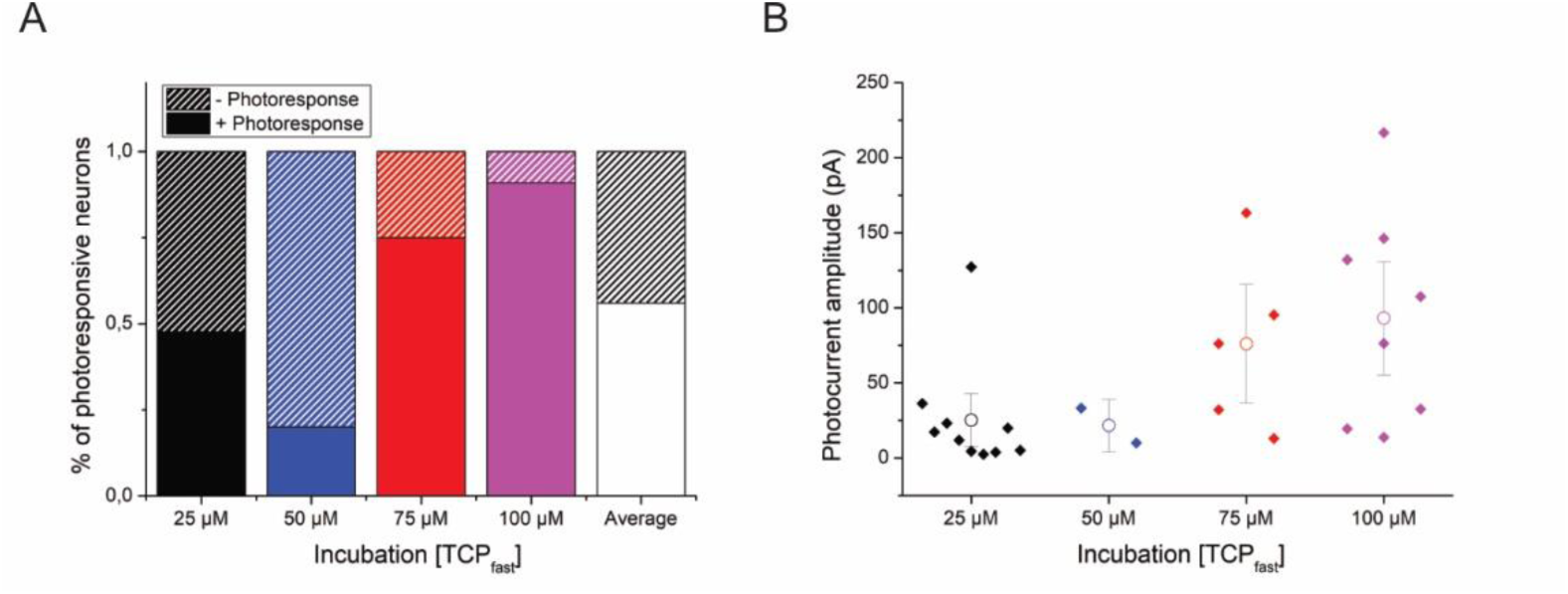
Optimization of TCP_fast_ conjugation conditions to endogenous neural receptors *in vitro*. **A.** Increasing TCP_fast_ concentration during incubation (2 min, pH9, 25-100 µM) leads to a higher fraction of cells displaying photoresponses. Fresh TCP_fast_ was obtained after 45 min head-tail click coupling reaction. **B.** Photocurrent amplitude increases proportionally to concentration of TCP_fast_ incubated. Note: From total num cells responding to perfusion of 300 µM glutamate, 56% had light response. Per concentration, 47,62% of cells incubated at 25 µM had light response (n=21); 20% of cells incubated at 50 µM had light response (n=10); 75% of cells incubated at 75 µM had light response (n=8); 90% of cells incubated at 100 µM had light response (n=11).

**Supplementary Figure S16.**
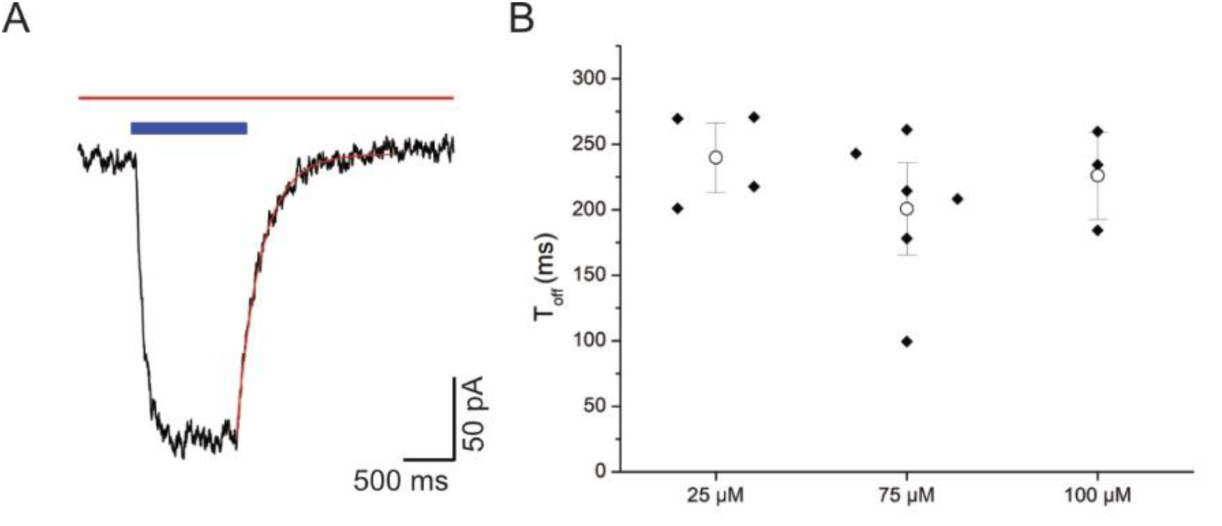
Fast relaxation lifetime of TCP_fast_ enables single wavelength control of photocurrents in hippocampal neurons. **A.** Representative current recording in response to 1 s blue light stimulation (blue bar) in presence of glutamate (300 μM, red bar) in bath solution. Whole cell voltage clamp mode recording in rat hippocampal neuron maintained 15 days in culture and incubated with TCP_fast_ (100 μM for 2 min at pH9). Exponential fit of the photocurrent to obtain T_off_ is showed in red. **B.** T_off_ average values obtained in neurons incubated with TCP_fast_ for 2 min at pH 9 at 25 μM (n=4), 75 μM (n=6) and 100 μM (n=3). Each data point is the average of T_off_ values obtained from fitting 5 different light pulses in the same cell. White dot is mean ± SE. The relaxation lifetime averaged for all concentrations is 220 ± 48 ms.

**Supplementary Figure S17.**
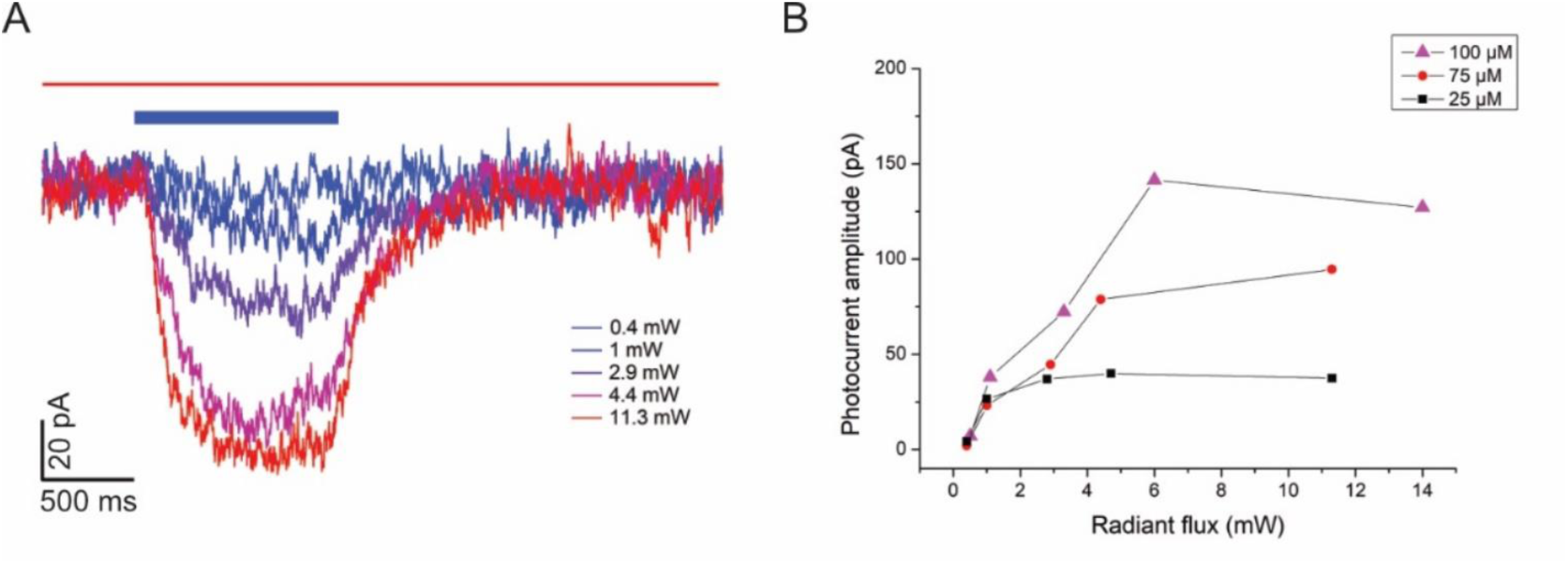
Photocurrent amplitude depends on light intensity. **A.** Superimposed representative photocurrent responses to different light intensities (473 nm, 1 s) in the same neuron. Power values are indicated in the label. Whole cell voltage clamp mode recording in dissociated rat hippocampal neurons maintained 12 days in culture and incubated with TCP_fast_ (75 μM for 2 min at pH9). **B.** Photocurrent amplitude as a function of radiant flux of different neurons incubated at different concentrations of TCP_fast_ for 2 min at pH9.

**Supplementary Figure S18.**
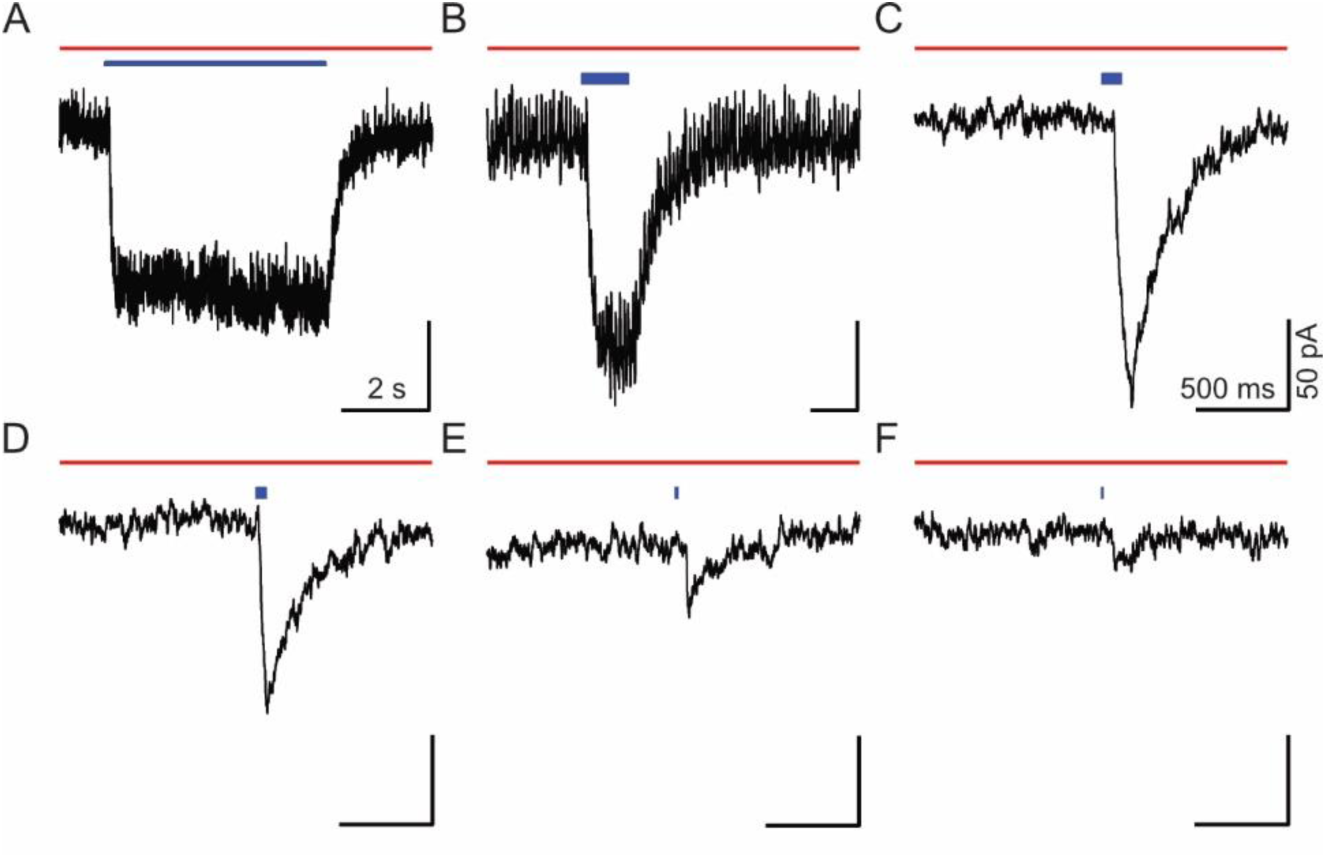
Neurons incubated with TCP_fast_ display reversible *cis*-on photocurrents in response to blue light pulses as short as a few milliseconds. Current recording in whole cell voltage clamp mode of dissociated rat hippocampal neurons maintained 15 days in culture and incubated with TCP_fast_ (100 μM for 2 min at pH9). Photocurrents elicited by illumination at λ_ex_ = 473 nm and duration 5 s (A), 500 ms (B), 100 ms (C), 50 ms (D), 10 ms (E) and 3 ms (F) in the presence of 300 µM glutamate (red bar).

**Supplementary Figure S19.**
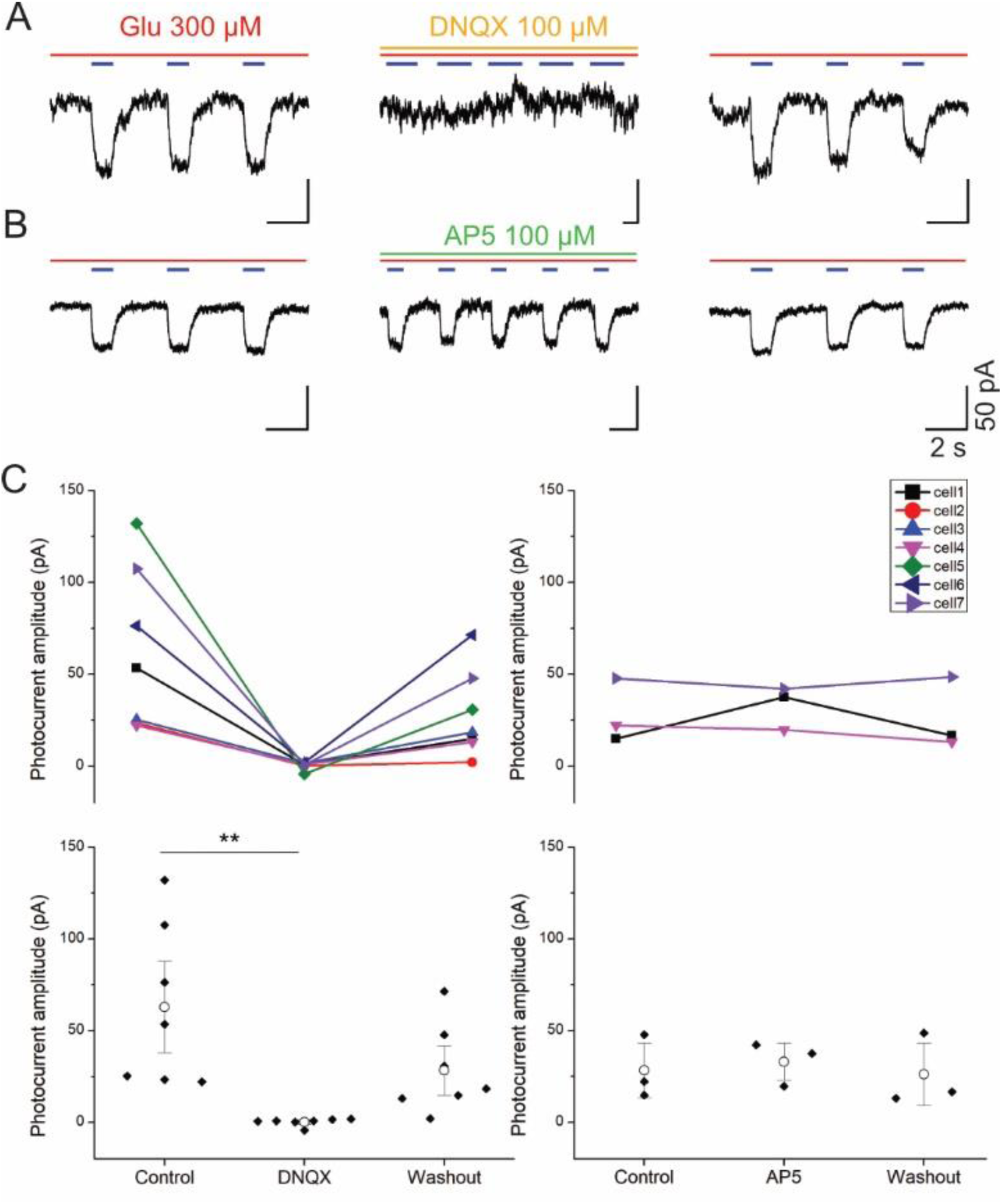
TCP_fast_ photocurrents in neurons are mediated by endogenous non-NMDA glutamate receptors. **A-B.** Current recordings in whole cell voltage clamp mode in rat hippocampal neurons maintained 15 days in culture and incubated with TCP_fast_ (100 μM for 2 min at pH9). Example traces of photocurrents elicited by irradiation at λ_ex_ = 473 nm (1 s, blue bars) in the presence of 300 µM glutamate (red bar). Photocurrents are reversibly blocked by perfusion of 100 µM DNQX (A, orange bar) and not affected by 100 µM AP5 (B, green bar). After washout and reperfusion of 300 µM glutamate photocurrents are recovered. **C.** Quantification of the effect of DNQX (100 µM, n = 7, p-value = 0.008) and AP5 (100 µM, n = 3, p-value = 0.56) on the photocurrent amplitude obtained from hippocampal neurons incubated with TCP_fast_ (25-75-100 μM for 2 min at pH9). Control and wash-out measurements were obtained after bath solution and glutamate perfusion. White dots indicate mean ± SE. Note: p-values obtained after performing Friedman test: nonparametric, data from any distribution; small samples; related samples.

## 7. *In vivo* photocontrol of neural activity in gerbil’s cochlea

**Supplementary Figure S20.**
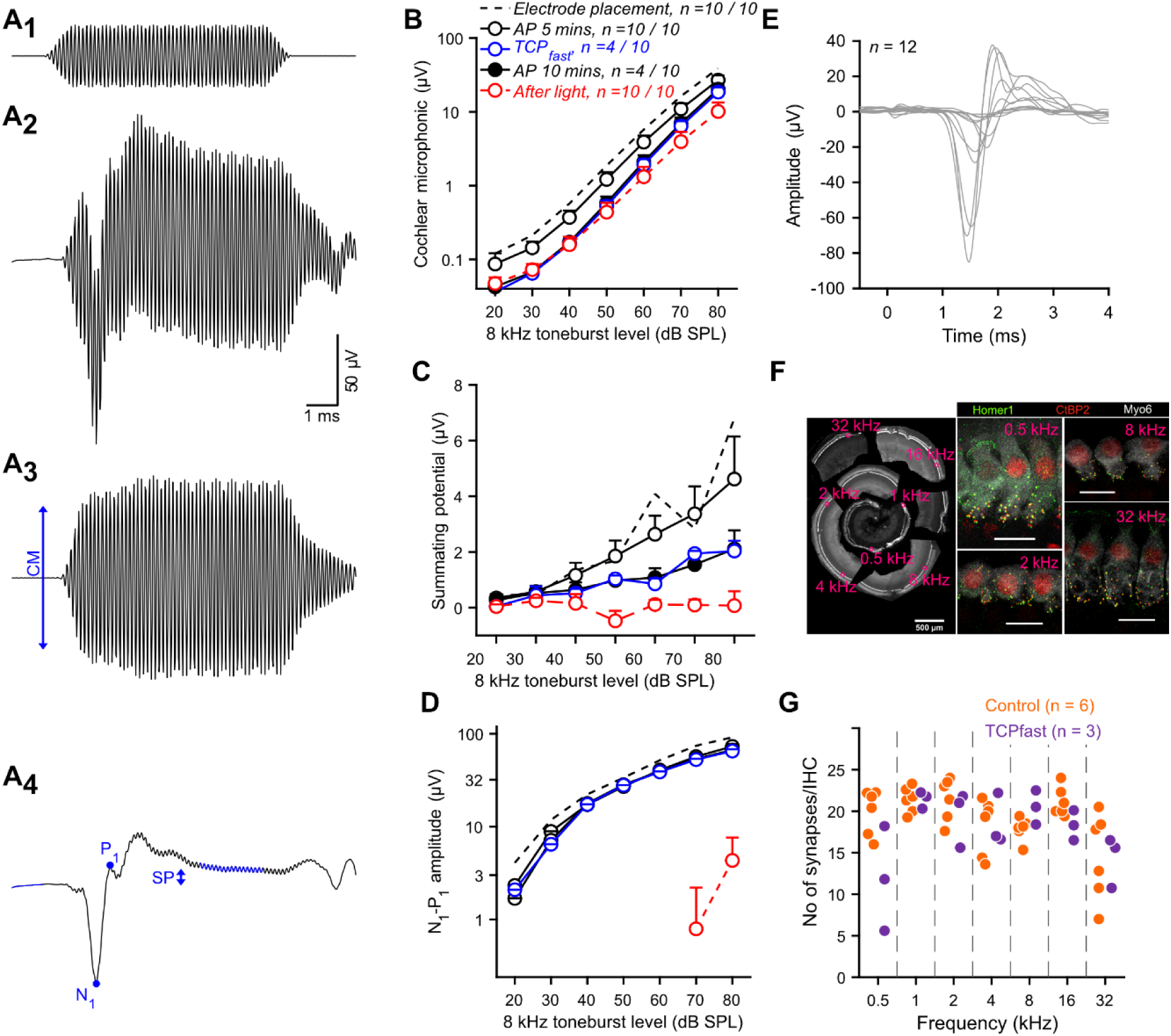
12.5 µM TCP_fast_ administration into the cochlea enables a transient optically evoked response of the SGNs followed by a loss of function of the inner hair cells and spiral ganglion neurons. **A.** Cochlear mass potentials (A_2_), recorded by a silver ball electrode implanted into the cochlear round window niche, in response to acoustic toneburst of 8 kHz (A_1_). The cochlear microphonic (CM, A_3_), reflecting outer hair cells activity, is extracted from the mass potential by a band passed filter centered on the stimulation frequency. The summating potential (SP, A_4_), reflecting inner hair cell activity, is measured as the difference between the base line and the plateau potential of the low-passed (< 3.5 kHz) filtered mass potential. The compound action potential (CAP, A_4_), reflecting the synchronous SGN first spike evoked by the sound stimulation, is measured as the difference between the negative peak N_1_ and the positive peak P_1_. **B-D.** Quantification (mean ± SEM) of the cochlear microphonic (B), summating potential (C) and CAP (D) as a function of the stimulation level after electrode placement (dashed line, *n* = 10), 5 min after artificial perilymph (AP) application defining the base line (open black circle, *n* = 10), following 12.5 µM TCP_fast_ application (open blue circle, *n* = 4), 10 min rinsing with AP (close dark circle, *n* = 4) and following light stimulation (red open circle, *n* = 10). **E.** Transient optically evoked CAPs (oCAP) recorded from 12 treated cochleae with 12.5 µM TCP_fast_. **F.** Right: Maximum projections of confocal stacks of immunolabelled gerbil IHC afferent synapses (IHC, anti-Myo6, gray; pre-synapse, anti-CtBP2/RIBEYE, red; post-synapse, anti-Homer1, green) at different tonotopic location (scale bar = 10 µm). Left: Montage of the low-magnification view of fragments of the full gerbil organ of Corti (scale bar = 500 µm). **G.** Quantification of the number of synapses per IHC between non-treated (*n* = 6) and 12.5 µM TCP_fast_ treated (*n* = 3) cochleae at different tonotopic regions (0.5, 1, 2, 4, 8, 16 and 32 kHz).

**Supplementary Figure S21.**
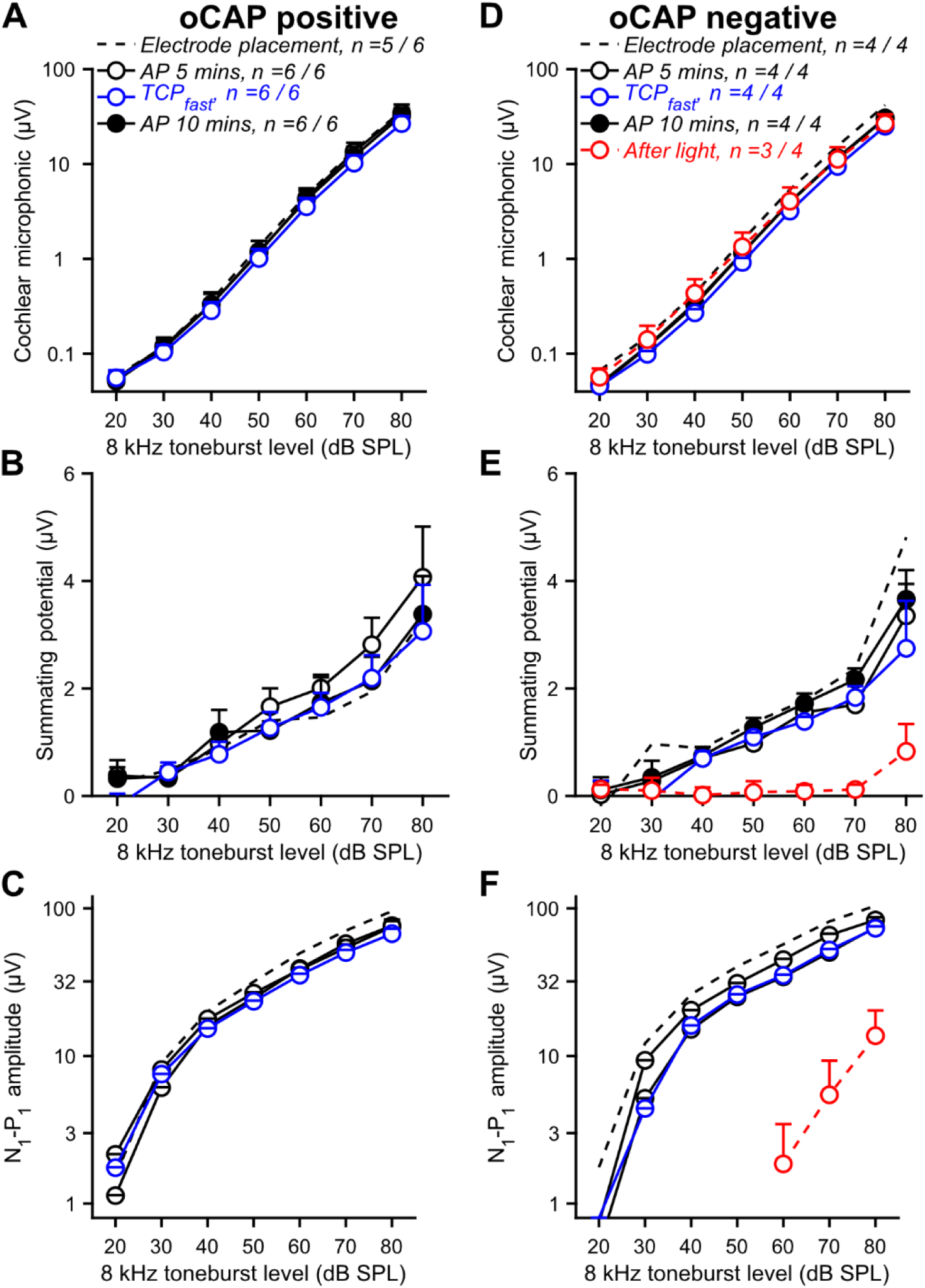
In 60% of the cases 2.5 µM TCP_fast_ application into the cochlea allows stable photoresponse in absence of toxicity for the organ of Corti. A-F. Quantification (mean ± SEM) of the cochlear microphonic (A,D), summating potential (B,E) and CAP (C,F) as a function of the stimulation level after electrode placement (dashed line), 5 min after artificial perilymph (AP) application defining the base line (open black circle), following 12.5 µM TCPfast application (open blue circle), 10 min rinsing with AP (close dark circle) and following light stimulation (red open circle) for the cochleae from which stable optically evoked CAPs were recorded (oCAP positive) or not (oCAP negative).

**Supplementary Figure S22.**
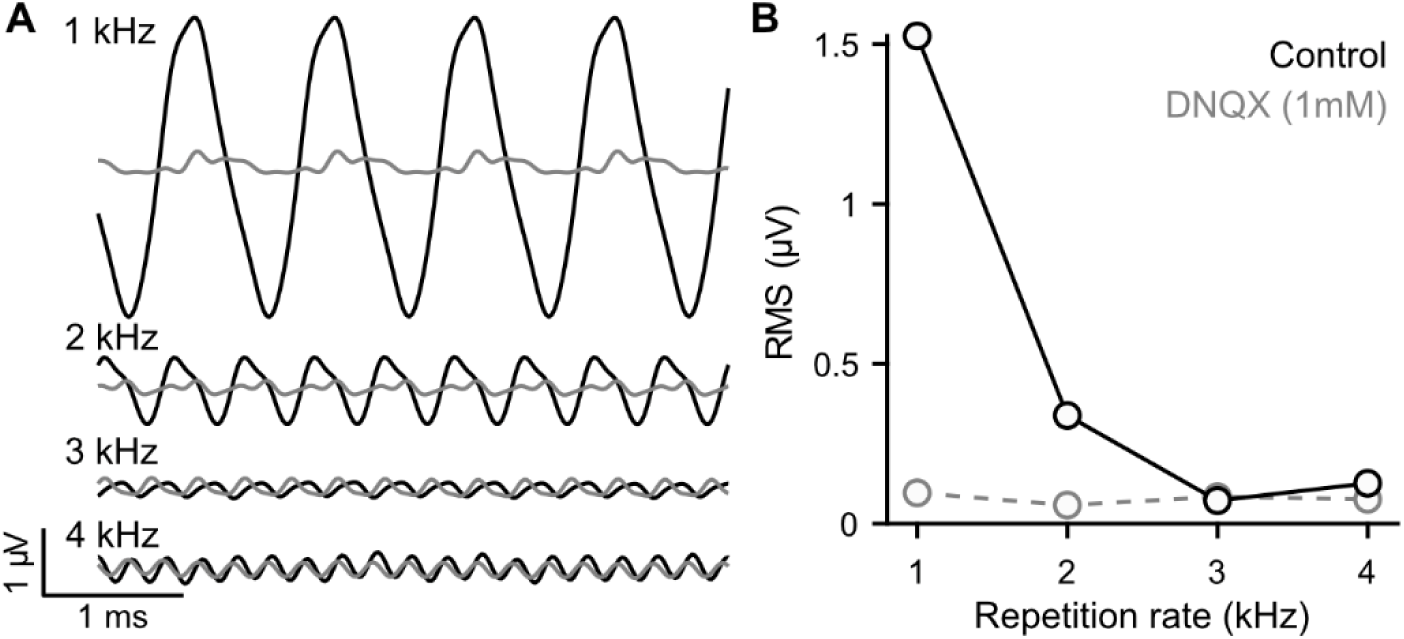
TCP_fast_ enables AMPA-mediated oCAPs up to stimulation rate of 2 kHz. **A.** oCAPs recorded from one cochlea in response to 1, 2, 3 and 4 kHz repetition rate before (black) and after DNQX (1 mM, gray). **B.** Measure of the oCAP amplitude (illustrated in A) as a function of the repetition rate before (black) and after DNQX application (gray).

